# Impaired development of neocortical circuits contributes to the neurological alterations in *DYRK1A* haploinsufficiency syndrome

**DOI:** 10.1101/438861

**Authors:** Juan Arranz, Elisa Balducci, Krisztina Arató, Gentzane Sánchez-Elexpuru, Sònia Najas, Alberto Parras, Elena Rebollo, Isabel Pijuan, Ionas Erb, Gaetano Verde, Ignasi Sahun, Maria J. Barallobre, José J. Lucas, Marina p. Sánchez, Susana de la Luna, Maria L. Arbonés

## Abstract

Autism spectrum disorders are early onset neurodevelopmental disorders characterized by deficits in social communication and restricted repetitive behaviors, yet they are quite heterogeneous in terms of their genetic basis and phenotypic manifestations. Recently, *de novo* pathogenic mutations in *DYRK1A*, a chromosome 21 gene associated to neuropathological traits of Down syndrome, have been identified in patients presenting a recognizable syndrome included in the autism spectrum. These mutations produce DYRK1A kinases with partial or complete absence of the catalytic domain, or they represent missense mutations located within this domain. Here, we undertook an extensive biochemical characterization of the *DYRK1A* missense mutations reported to date and show that most of them, but not all, result in enzymatically dead DYRK1A proteins. We also show that haploinsufficient *Dyrk1a*^+/-^ mutant mice mirror the neurological traits associated with the human pathology, such as defective social interactions, stereotypic behaviors and epileptic activity. These mutant mice present altered proportions of excitatory and inhibitory neocortical neurons and synapses. Moreover, we provide evidence that alterations in the production of cortical excitatory neurons are contributing to these defects. Indeed, by the end of the neurogenic period, the expression of developmental regulated genes involved in neuron differentiation and/or activity is altered. Therefore, our data indicate that altered neocortical neurogenesis could critically affect the formation of cortical circuits, thereby contributing to the neuropathological changes in *DYRK1A* haploinsufficiency syndrome.

## 1. Introduction

Autism spectrum disorder (ASD) is a neurodevelopmental disorder characterized by social communication deficits and restricted repetitive behaviors (Lord and Bishop, 2015), and it is frequent associated with intellectual disability (ID), language deficits and seizures (Geschwind, 2009; Sztainberg and Zoghbi, 2016). The causes of ASD remain largely unknown, although a genetic cause has been identified in around 25% of cases (Huguet et al., 2013). These genetic alterations include chromosomal rearrangements, *de novo* copy-number variants and *de novo* mutations in the coding-sequence of genes associated with chromatin remodeling, mRNA translation or synaptic function (de la Torre-Ubieta et al., 2016). Additionally, alterations in the production or migration of neocortical neurons are a point of convergence for ASD and ID (de la Torre-Ubieta et al., 2016; Ernst, 2016; Packer, 2016).

DYRK1A is a member of the conserved dual-specificity tyrosine-regulated kinase (DYRK) family of protein kinases (Aranda et al., 2011) that has different functions in the nervous system (Tejedor and Hammerle, 2011). This kinase influences brain growth, an activity that is conserved across evolution (Fotaki et al., 2002; Kim et al., 2017; Tejedor et al., 1995). *DYRK1A* is located within the Down syndrome (DS) critical region on human chromosome 21 (Guimera et al., 1996). There is evidence that triplication of the gene contributes to neurogenic cortical defects (Najas et al., 2015) and other neurological deficits in DS, making it a potential drug target for DS-associated neuropathologies (Becker et al., 2014). Recently, mutations in *DYRK1A* have been identified in a recognizable syndromic disorder (*DYRK1A* haploinsufficiency syndrome [DHS], also known as DYRK1A-related intellectual disability syndrome; ORPHANET:464306; OMIM:614104). ASD-related deficits are common clinical manifestations in DHS, which include moderate to severe ID, intrauterine growth retardation, developmental delay, microcephaly, seizures, speech problems, motor gait disturbances and a dysmorphic *facies* (Earl et al., 2017; Luco et al., 2016; van Bon et al., 2016). The mutations identified to date in patients with DHS are *de novo*, involving chromosomal rearrangements (Courcet et al., 2012; Moller et al., 2008; van Bon et al., 2011), small insertions or deletions, and nonsense mutations (Courcet et al., 2012; Earl et al., 2017; O’Roak et al., 2012; van Bon et al., 2016 and references therein). These mutations lead to the production of truncated DYRK1A proteins that lack partially o totally the kinase domain and that are therefore catalytically inactive. *DYRK1A* missense mutations have also been identified in patients with a distinctive DHS phenotype (Bronicki et al., 2015; Dang et al., 2018; De Rubeis et al., 2014; Deciphering Developmental Disorders, 2015; Evers et al., 2017; Ji et al., 2015; Ruaud et al., 2015; Stessman et al., 2017; Trujillano et al., 2017; Wang et al., 2016; Zhang et al., 2015). The structural modeling of these mutations predicts that they are loss-of-function (LoF) mutations (Evers et al., 2017; Ji et al., 2015). However, experimental data supporting this prediction have been reported only for a few of them (Dang et al., 2018; Widowati et al., 2018).

The *Dyrk1a^+/-^* mouse displays some of the core traits of DHS, including developmental delay, microcephaly, gait disturbances and learning problems (Arque et al., 2008; Fotaki et al., 2002; Fotaki et al., 2004). The cell density in the neocortex of the adult mutant mouse is altered (Fotaki et al., 2002; Guedj et al., 2012) but the origin of this alteration and the neuronal components affected are unknown. To gain additional insight into the pathogenesis of DHS, we have performed a biochemical study to assess the impact of all reported missense mutations within the catalytic domain on *DYRK1A* activity. Furthermore, we analyzed the cytoarchitecture and gene expression profile of the neocortex in *Dyrk1a^+/-^* mice, also performing behavioral tests and obtaining electroencephalogram (EEG) recordings from these animals. Our results indicate that defects in the production of excitatory neocortical neurons are critical to the neuropathology of DHS.

## 2. Materials and methods

### 2.1. Plasmids

The plasmids to express HA-tagged DYRK1A have been described previously (Alvarez et al., 2003: human isoform 754 aa; NP_569120), both the wild type (WT) or a kinase inactive version in which the ATP binding lysine 179 (K188R in the 763 aa isoform) was replaced by arginine. Single nucleotide mutations (numbering based on the 763 aa isoform, as appears in most publications: Supplementary Table 1 and Fig. 1A) were introduced into the HA-DYRK1A expression plasmid using the QuickChange Multi Site-Directed Mutagenesis Kit, according to the manufacturer’s instructions (Agilent Technologies) and with primers specific to each mutant (Supplementary Table 2). The A469fs* and A489fs* mutations were produced by a T or C deletion, respectively, and the same alterations were created in the DYRK1A expression plasmid. The plasmids generated by site-directed mutagenesis were verified by DNA sequencing and the transfection efficiency was assessed by co-transfection with a green fluorescent protein (GFP) expression plasmid (pEGFP-C1, Clontech).

**Fig. 1.**
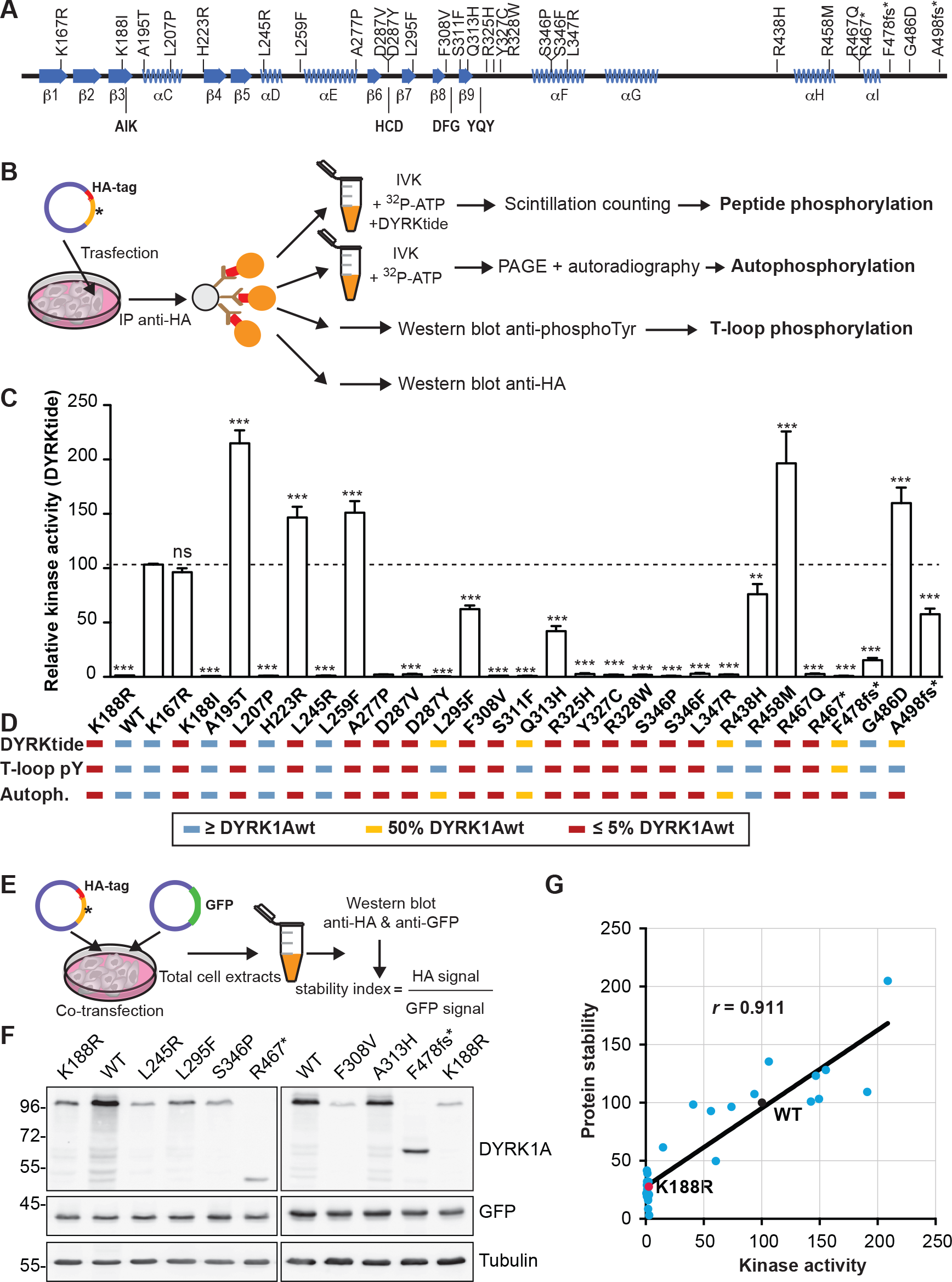
*DYRK1A* missense mutations affect DYRK1A kinase activity, auto-phosphorylation and protein stability. (A) Representation of the secondary protein structure of the DYRK1A catalytic domain, indicating the location of the mutants used in this study: AIK, HCD, DFG and YQY correspond to key functional elements (Kannan and Neuwald, 2004). (B) Experimental procedure followed to analyze the parameters summarized in (C) and (D). (C) The graph represents the ability of the mutants to phosphorylate the DYRKtide peptide, with the WT kinase activity arbitrarily set as 100. The catalytically inactive mutant K188R was also included in the assay (n=3 independent experiments; \*\*\**p* < 0.001, ns=not significant, unpaired 2-tailed Mann-Whitney’s test). (D) Summary of the mutants’ activity measured as the substrate phosphorylation, auto-phosphorylation and T-loop auto-phosphorylation (see Supplementary Fig. 1B and C). (E, F) Scheme of the assay used to assess the impact of the mutations on protein accumulation (E). A representative experiment is shown (F; see also Supplementary Fig. 2 for quantification). (G) Correlation analysis of the activity and stability of the DYRK1A mutants. The WT protein and the kinase-inactive K188R mutant are indicated as black and red dots, respectively (Pearson’s correlation, *r* = 0.9211; *p* < 0.0001).

### 2.2. Animals

We used embryos and postnatal and adult *Dyrk1a*^+/-^ mice and their control *Dyrk1a*^+/+^ littermates, generated and genotyped as described elsewhere (Fotaki et al., 2002; Najas et al., 2015). The day of the vaginal plug was defined as E0 and the day of birth was defined as P0. After weaning, mice from the same litter and of the same gender were housed in groups. The animals were maintained at the PCB-PRBB Animal Facility in ventilated cages on a 12 h light/dark cycle, at approximately 20°C and in 60% humidity, and with food and water supplied *ad libitum*. For bromodeoxyuridine (BrdU) birthdating experiments, pregnant females were peritoneally injected with a BrdU solution (50 mg/kg; Sigma), two injections spaced 4 h, and pups were collected and processed at P7. Experimental procedures were carried out in accordance with the European Union guidelines (Directive 2010/63/EU) and the protocols were approved by the Ethics Committee of the CSIC, PCB-PRBB, CBM and Fundación Jiménez Díaz.

### 2.3. Cell culture and transfection

The HEK293T cell line was obtained from the American Type Culture Collection (www.atcc.org) and used for the exogenous expression of DYRK1A mutants. The cells were maintained at 37°C in Dulbecco’s modified Eagle medium (DMEM; Invitrogen), with 10% fetal bovine serum (FBS; Invitrogen) and supplemented with antibiotics (100 μg/ml penicillin and 100 μg/ml streptomycin; Invitrogen). The cells were transfected by the calcium phosphate precipitation method and processed 48 h after transfection.

### 2.4. *In vitro* kinase (IVK) assays

Cells were washed in phosphate buffered saline (PBS) and then lysed in HEPES lysis buffer (50 mM Hepes [pH 7.4], 150 mM NaCl, 2 mM EDTA, 1% NP-40) supplemented with a protease inhibitor cocktail (#11836170001, Roche Life Science), 30 mM sodium pyrophosphate, 25 mM NaF and 2 mM sodium orthovanadate. The lysates were cleared by centrifugation and incubated overnight at 4°C with protein G-conjugated magnetic beads (Dynabeads, Invitrogen) previously bound to an antibody against HA (Covance, #MMS-101R). The beads were then washed 3 times with HEPES lysis buffer and used for either IVK assays or to probe Western blots to control for the presence of HA-tagged DYRK1A. For the IVK assays, immunocomplexes were washed in kinase buffer (25 mM HEPES [pH 7.4], 5 mM MgCl_2_, 5 mM MnCl_2_, 0.5 mM DTT) and further incubated for 20 min at 30°C in 20 μl of kinase buffer containing 50 μM ATP, [λ^32^P]-ATP (2.5×10^-3^ μCi/pmol) and with 200 μM DYRKtide as the substrate peptide. The incorporation of ^32^P was determined as described previously (Himpel et al., 2000), and the kinase activity was normalized to the amount of DYRK1A protein present in the immunocomplexes determined in Western blots (see Supplementary Materials and Methods for details and Table 3 for information on primary antibodies). DYRK1A autophosphorylation was analyzed by SDS-PAGE fractionation of the immunocomplexes and exposure of the dried gel to film.

### 2.5. RNA extraction, microarray analysis and RT-qPCR

Postnatal day (P) 0 and P7 mice were sacrificed by decapitation and the brain cortices were dissected out and stored at -80°C. The tissue samples were homogenized in a Polytron and the total RNA was extracted with the TriPure Isolation Reagent (Roche). A RNA clean-up step was performed using the RNeasy Mini Kit (Qiagen) followed by DNAse I treatment (Ambion). Only RNA samples with an integrity number above 8.0 (Agilent 2001 Bioanalyzer) were used for further analysis. For microarray studies, the total RNA was hybridized to an Affymetrix Mouse GeneChip 430 2.0 Array. RNA was analyzed by reverse transcription coupled to quantitative PCR (the sequence of the qPCR primers is found in Supplementary Table 4) or using a low-density array (probes included in Supplementary Table 5). For more detailed information see Supplementary Materials and Methods.

### 2.6. Video-EEG recordings

Video-EEG recordings of adult *Dyrk1a*^+/+^ and *Dyrk1a*^+/-^ male mice (8 animals each genotype) aged 5-6 months were obtained as described elsewhere (Garcia-Cabrero et al., 2012). Two home-made EEG stainless steel electrodes were implanted symmetrically into an anesthetized mouse over the cortex in front of bregma, and the ground and reference electrodes were placed posterior to lambda (Fig. 2C). A plastic pedestal (Plastic One Inc., Roanoke, VA, USA) was used to attach the pins of the electrodes, and the headset was fixed to the skull and the wound closed with dental cement (Selectaplus CN, Dentsply DeTrey GmbH, Dreireich, Germany). Four days after surgery, synchronized video-EEG activity was recorded for 48 h in freely moving mice using a computer-based system (Natus Neurowork EEG, Natus Medical Inc., San Carlos, CA, USA). EEG seizure activity was examined by inspecting the entire EEG records and behavioral correlations were reviewed within the corresponding video-type segments.

**Fig. 2.**
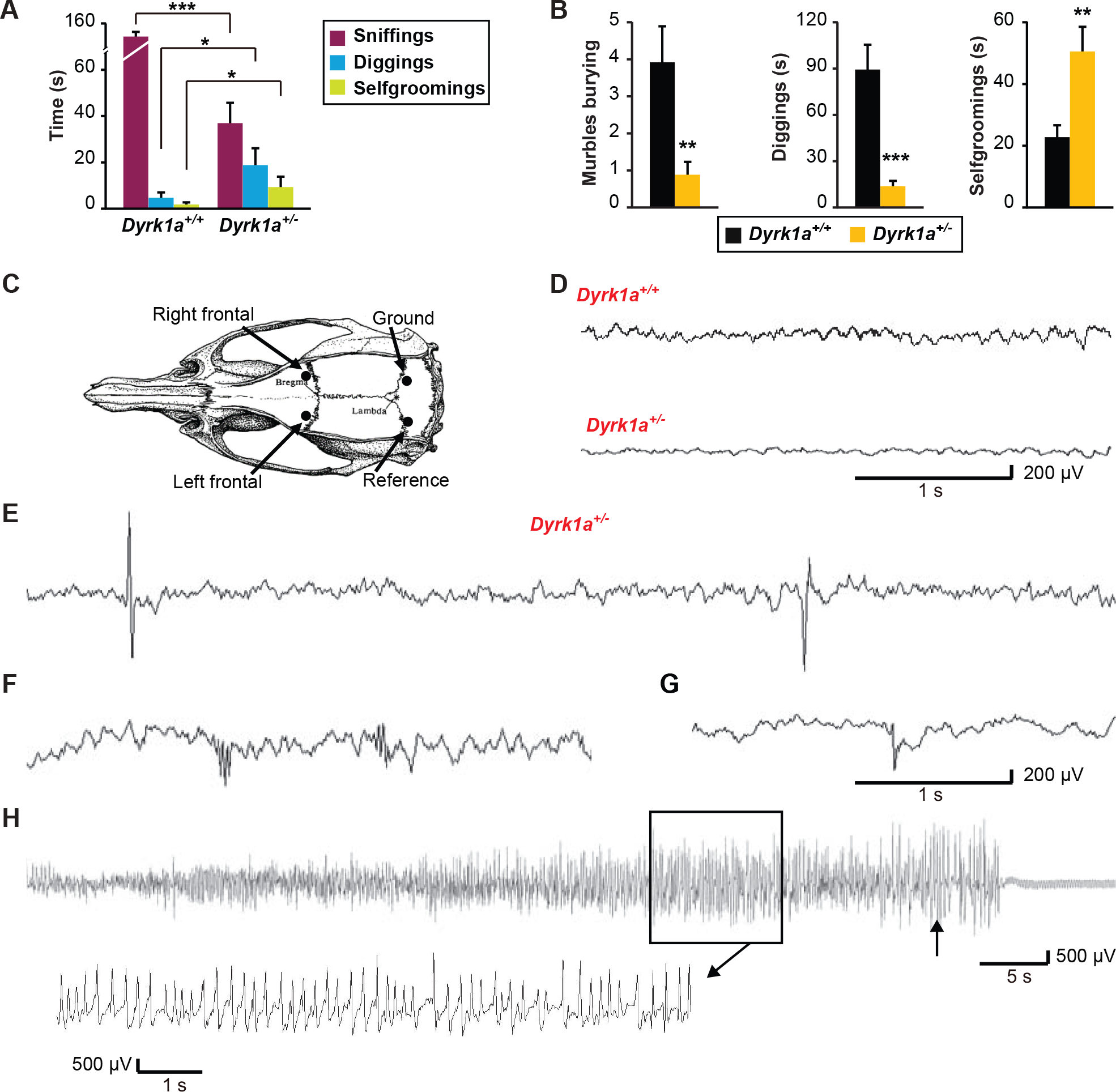
*Dyrk1a*^+/-^ mice display social deficits, repetitive behavior and epileptic activity. (A) Social interaction test. Time spent by adult mice on social interactions (sniffing) and repetitive behavior (digging and self-grooming). (B) Marble-burying test in which the number of buried marbles, and the time spent by adult mice digging and on self-grooming was recorded over 20 min (n=12-15 mice each genotype: \**p* < 0.05, \*\**p* < 0.01, \*\*\**p* < 0.001, Student’s *t*-test). (C) Schematic drawing of a mouse skull indicating the electrode placement for EEG recordings. (D) Fragments of representative EEG recordings of the basal activity in *Dyrk1a*^+/+^ and *Dyrk1a*^+/-^ mice. (E-G) Representative EEG recordings of *Dyrk1a*^+/-^ mice showing the spikes (E), polyspikes (F) and spike-wave discharges (G). (H) EEG recording of a spontaneous generalized tonic-clonic seizure lasting 74 s in a *Dyrk1a*^+/-^ mouse (see also Supplementary Movie), with a 10 s amplified portion (black square) displaying spike-wave activity at a frequency of 4-5 Hz. The arrow indicates the beginning of the clonic phase.

### 2.7. Behavioral Tests

The social and stereotyped behaviors (social interaction and marble-burying tests) of adult 4-6 month old *Dyrk1a*^+/+^ and *Dyrk1a*^+/-^ male mice (12-15 animals each genotype) were tested in a dedicated room for behavioral studies during the light phase. Before the experiment, the animals were habituated to the room for at least 30 min and during the test, the mice were recorded from above with a video-camera CCTV (Sonny Corporation). The animal’s behavior was analyzed by an observer blind to their genotype using the SMART 3.0 software (Panlab Harvard Apparatus). See Supplementary Materials and Methods for more details on these procedures as well as for the UsV analysis.

### 2.8. Tissue preparation, immunostaining, and cell and synapse counts

To obtain brain tissue, P7 and adult mice were deeply anesthetized and transcardially perfused with 4% paraformaldehyde. The mouse’s brain was removed and post-fixed, and cryotome or vibratome brain sections were immunostained as described previously (Barallobre et al., 2014). To quantify neurons and synapses, images were obtained on a Leica AF6500 motorized wide-field microscope and a Zeiss LSM780 confocal microscope, respectively. To quantify BrdU-labeled cells, images were obtained on an Olympus BX51 motorized microscope with a JVC digital color camera. The procedures for immunostaining, and cell and synapse quantification are indicated in the Supplementary Materials and Methods. The source of the primary antibodies for these procedures is indicated in Supplementary Table 6.

### 2.9. Statistical analysis

Statistical analyses were performed with GraphPad Prism v5.0a (GraphPad Software) or with SPSS (IBM Analytics Software). The data from the IVK studies were analyzed with a 2-tailed unpaired Mann-Whitney test, synapse counts with a nested one-way ANOVA test, and the rest of the analysis was performed with a two-tailed unpaired Student’s *t*-test. Differences were considered significant at *p* < 0.05. In the histograms, the data are represented as the mean ± standard error of the mean (SEM).

## 3. Results

### 3.1 *DYRK1A* missense mutations in individuals with DYRK1A haploinsufficiency are loss-of-function mutations

The kinase activity associated with most of *DYRK1A* missense mutations has been inferred from their position within the DYRK1A catalytic domain (Evers et al., 2017). Given that experimental data supporting the predicted models have been reported very recently for only a few *DYRK1A* missense mutations (Widowati et al., 2018), we set out to analyze the kinase activity of all missense mutations within the DYRK1A catalytic domain published to date and some others included in the ClinVar database (www.ncbi.nlm.nih.gov/clinvar/?term=DYRK1A[gene]). We also included 2 missense mutations in the non-catalytic C-terminal region (R528W and T588N) and 3 mutations generating truncated proteins at the end of the kinase domain and predicted to contain the whole catalytic domain (Fig. 1A and Supplementary Table 1). These mutations were introduced into a DYRK1A expression plasmid and the catalytic activity of each of the mutants was evaluated using an IVK assay with the DYRKtide peptide as the substrate (see experimental design in Fig. 1B). The activity of most of the missense mutants was comparable to that of the DYRK1A K188R kinase-dead mutant (Fig. 1C), although the L295F, Q313H and R438H mutants rendered proteins with half the activity of the WT protein (Fig. 1C). However, some of the mutations did not alter the DYRK1A catalytic activity (K167R and T588N), and six even enhanced the enzyme’s kinase activity: A195T, H223R, L259F, R458M, G486D and R528W (Fig. 1C and Supplementary Fig. 1A). Notably, the activity of the truncated proteins increased with the length of the resulting protein, with truncations at R467 (before the α-helix I) devoid of activity, and at F478 (downstream α-helix I) with just 10% activity (Fig. 1C). Moreover, the fact that the A498Pfs93* mutant does not have similar catalytic activity to the WT protein might indicate that all or part of the non-catalytic C-terminal region is required for full DYRK1A kinase activity. Of note, the crystal structures of DYRK1A have been obtained with truncated proteins that, based on our data, are not fully active (Soundararajan et al., 2013).

Considering the mode of activation of DYRK1A (Lochhead et al., 2005), the negative results in the IVK assays could be due to defective autophosphorylation of the Tyr residue in the activation loop, or to the inability of the mutants to phosphorylate an exogenous substrate. The lack of activity on the substrate was correlated with a lack of Tyr phosphorylation (Fig. 1D), suggesting that the mutations altered the activation of the kinase. We also evaluated the autophosphorylation of the mutant proteins (Fig. 1B), an activity associated with the mature kinase (Himpel et al., 2001), and only the catalytically active mutant enzymes were seen to autophosphorylate, with the exception of the truncated mutant A498Pfs93* (Fig. 1D and Supplementary Fig. 1B and C). In this latter case, either the autophosphorylation sites are located in the missing C-terminal region or the mutation alters the substrate preference of the protein. Together, these results indicate that most but not all of the missense DYRK1A mutants reported to date in DHS are *bona-fide* LoF mutants.

There is deficient accumulation of the kinase-dead DYRK1A K188R mutant (Kii et al., 2016) and therefore, we evaluated the quantitative expression of these mutants and their stability (Fig. 1E: stability index). We confirmed the reduced accumulation of the K188R mutant relative to the WT protein (Fig. 1F), and a similar behavior was observed for other catalytically inactive mutants (Fig. 1F and Supplementary Fig. 2). In fact, kinase activity and protein stability were positively correlated (Fig. 1G), suggesting that in heterozygosis, the proteins expressed from the mutant allele will not accumulate as much as the WT protein, thereby reducing their potential dominant-negative activity.

### 3.2. Stereotyped behavior and epileptic activity of *Dyrk1a*^+/-^ mice

Seizures, stereotypies and social anxiety are frequent in DHS (Earl et al., 2017; van Bon et al., 2016). The *Dyrk1a*^+/-^ mouse is more sensitive to the convulsive agent pentylenetetrazol (Souchet et al., 2014) and it displays anxiety-like behavior in the open field (Fotaki et al., 2004), although ASD-like behaviors and epileptiform activity have not been reported in this model. Recently, a mouse carrying a frame-shift mutation in *Dyrk1a* has been generated and mice heterozygous for this mutation develop deficits in social contacts and communicative ultrasonic vocalization (UsV), as well as hyperthermia-induced seizures (Raveau et al., 2018). *Dyrk1a*^+/-^ pups displayed similar deficits in UsV when separated from their mothers (Supplementary Fig. 3A). Moreover, adult *Dyrk1a*^+/-^ mice engaged in shorter social olfactory interactions (oral-oral or oral-genital sniffing) in the resident-intruder test, while more time was spent on repetitive digging and self-grooming (Fig. 2A). Stereotyped behaviors were further evaluated in the marble-burying test, and the *Dyrk1a*^+/-^ mutant mice buried fewer marbles and spent less time digging than their WT littermates (Fig. 2B and Supplementary Fig. 3B). Furthermore, *Dyrk1a*^+/-^ mutants spent less time exploring the central region of the cage where the marbles were buried (*Dyrk1a*^+/+^ 288±41 s, *Dyrk1a*^+/-^ 133±15 s; *p* <0.001, Student’s *t*-test). This time difference is unlikely to be caused by less locomotor activity, as both genotypes entered the central region a similar number of times (*Dyrk1a*^+/+^ 41±5, *Dyrk1a*^+/-^ 35±4 s; *p* <0.314, Student’s *t*-test). Rather, *Dyrk1a*^+/-^ mice spent significantly longer on repetitive self-grooming (Fig. 2B). Together, these results show that in addition to deficits in social communication, haploinsufficient *Dyrk1a*^+/-^ mice develop repetitive and stereotyped behaviors.

The epileptic activity in *Dyrk1a*^+/-^ mice was evaluated from video-EGG recordings (Fig. 2C-H), which revealed differences in the basal activity between the distinct genotypes (*Dyrk1a*^+/+^ 9-12 Hz; *Dyrk1a*^+/-^ 6-7 Hz: Fig. 2D). Spontaneous myoclonic jerks that corresponded to interictal epileptiform activity were evident in *Dyrk1a*^+/-^ mice, with isolated spikes, polyspikes and spike-wave discharges (Fig. 2E-G). These mutant mice also experienced periods of immobility, during which grooming, feeding and exploration were suppressed. Such periods of inactivity, alternating with shorter ones of epileptic activity, were characterized by groups of spikes and polyspikes in the EEG recordings (Fig. 2E and F). At times, the epileptic activity in *Dyrk1a*^+/-^ mice coincided with situations of moderate handling stress. In half of the *Dyrk1a*^+/-^ mice (4 out of 8), spontaneous generalized tonic-clonic seizures were recorded that lasted for different periods, with distinct lengths of tonic and clonic phases, and motor correlates that corresponded to stage 5-6 in the modified Racine Scale (Luttjohann et al., 2009; see Fig. 2H and Supplementary Movie). Notably, about half of the patients with mutations in *DYRK1A* develop epilepsy, displaying atonic attacks, absences and generalized myoclonic seizures (van Bon et al., 2016). Thus, epileptic activity seems to be a common characteristic in humans and mice with DHS.

### 3.3. Alterations in neuron numbers in the developing neocortex of *Dyrk1a*^+/-^ mice

The neocortex is a six-layered structure essential for higher cognitive functions and sensory perception. Most neocortical neurons (more than 80% in the mouse) are projection excitatory neurons that extend their axons to intracortical or subcortical targets and use glutamate as neurotransmitter. The distinct types of excitatory neurons are produced in the embryo dorsal telencephalon (from E11.5 to E17.5 in the mouse) in overlapping temporal waves. The first neurons formed are those that are closest to the ventricle (layer VI neurons) and the last formed are those neurons located in the most superficial layers (layers II-III neurons) (Florio and Huttner, 2014; Molyneaux et al., 2007). We previously showed that production of the early-born cortical neurons is enhanced in *Dyrk1a*^+/-^ embryos (Najas et al., 2015). To obtain further evidence as to how *Dyrk1a* haploinsufficiency affects corticogenesis, we counted the number of neurons that express the neuronal marker NeuN in the internal (V-VI) and external (II-IV) layers of the somatosensorial cortex (SSC) at P7 (Fig. 3A), when radial migration has ended and excitatory neurons adopt their final positions (Miller, 1995). The neocortical layers were thinner in postnatal *Dyrk1a^+/-^* mice, while they contained more neurons than their *Dyrk1a*^+/+^ littermates (Fig. 3B and C). The fact that the neuronal densities in the internal and external layers of *Dyrk1a^+/-^* mutants are similarly affected (Fig. 3C) suggests that *Dyrk1a* haploinsufficiency may alter neuron production along the neurogenic phase of cortical development. This possibility was evaluated with BrdU-birthdating analysis (Fig. 3D-F). The results showed that P7 *Dyrk1a^+/-^* animals had more BrdU^+^ cells in the SSC than WT animals when the BrdU labeling was performed at E13.5 (Fig. 3D), indicating that neurogenesis is enhanced in the *Dyrk1a^+/-^* mutant at this developmental stage. By contrast, *Dyrk1a*^+/+^ and *Dyrk1a^+/-^* animals displayed similar numbers of BrdU^+^ cells when the treatment was performed at E15.5 (Fig. 3F). Additional BrdU-labeling experiments indicated that cortical neurogenesis in *Dyrk1a^+/-^* embryos ends at the correct time (Supplementary Fig. 5). Consistent with the temporal generation of cortical excitatory neurons (Molyneaux et al., 2007), the BrdU-labeled cells generated at E15.5 were concentrated in the external layers of P7 WT animals, while the labeled cells generated at E13.5 showed a broader layer distribution, with BrdU^+^ cells located in both internal and external layers (Fig. 3E and G). Additionally, the distribution of BrdU-labeled cells in *Dyrk1a^+/-^* animals was similar to that in WT animals (Fig. 3E and G), indicating that radial migration of excitatory neurons is not affected in the *Dyrk1a^+/-^* model. Taking all the results together, we conclude that the augmented neuron density in the neocortex of *Dyrk1a^+/-^* mutants is the result of an increased neuron production during early and mid-corticogenesis.

**Fig. 3.**
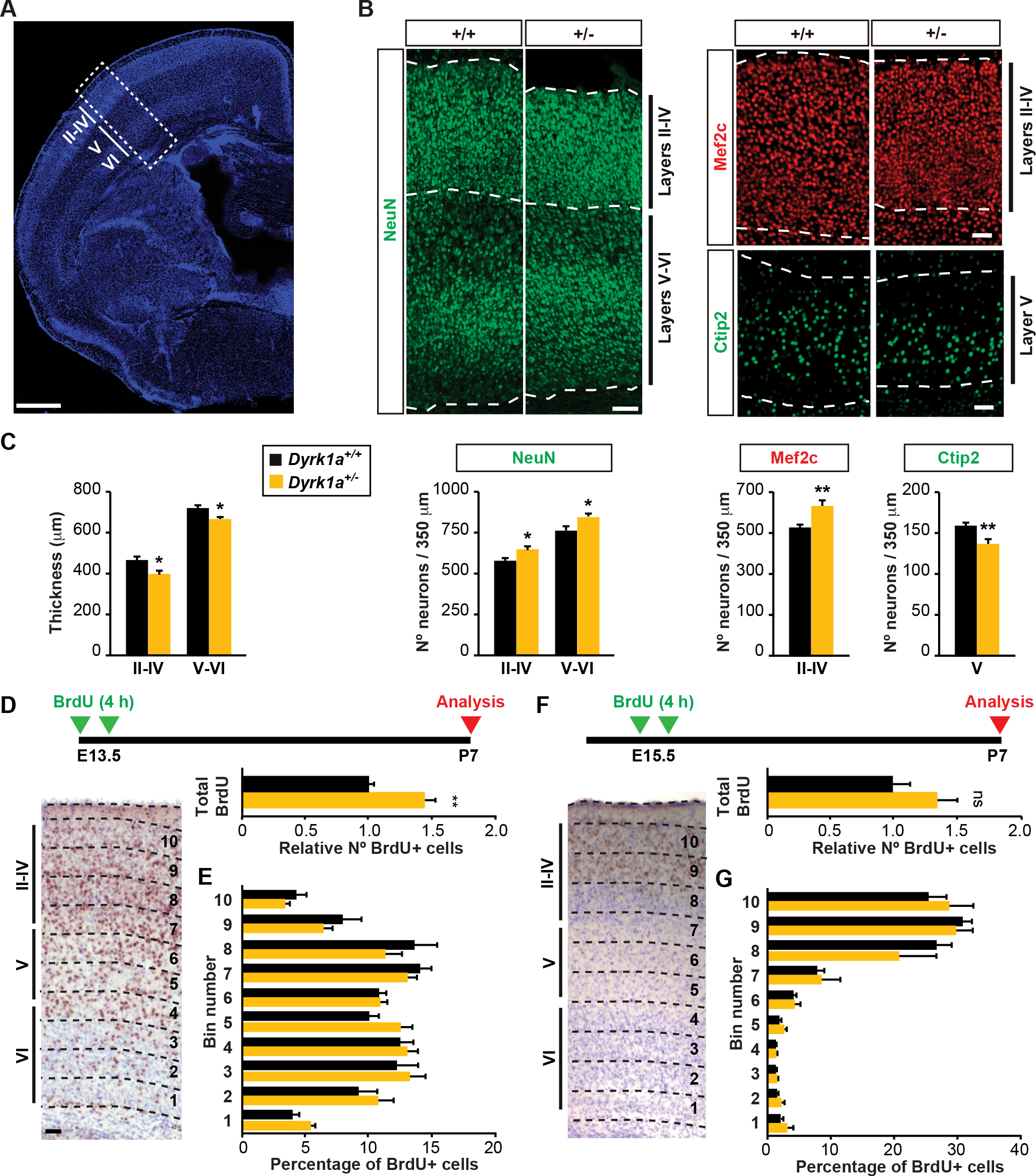
*Dyrk1a*^+/-^ mice show altered numbers of excitatory neurons in the postnatal neocortex. (A) Coronal brain section from a P7 *Dyrk1a*^+/+^ mouse stained with Hoechst indicating the position of the neocortical layers II to VI. Quantification in the somatosensorial cortex (SSC, white dashed rectangle). (B) Representative images of *Dyrk1a*^+/+^ (+/+) and *Dyrk1a*^+/-^ (+/-) sections immunolabeled for NeuN, Mef2c or Citp2. (C) Histograms showing the thickness and number of neurons positive for the markers indicated in a 350 μm wide column of the external (layers II-IV) and internal (layers V-VI) layers of the neocortex (n=4-5 animals each genotype). (D-G) Schedule of the BrdU-labeling protocol to estimate neuron production in E13.5 (D) and E15.5 (F) embryos and representative images of *Dyrk1a*^+/+^ sections immunolabeled for BrdU (brown signal) and the nuclei stained with Nissl. Histograms represent the total number of BrdU^+^ cells in *Dyrk1a*^+/-^ animals relative to that in *Dyrk1a*^+/+^ animals arbitrarily set as 1 (D-F), and the distribution of these cells in 10 equal bins, represented as the percentage of BrdU^+^ cells in each bin (E, G) (BrdU^+^ cells at E13.5, n=7-9 animals each genotype; BrdU^+^ cells at E15.5, n=5-6 animals each genotype). ns= not significant, \**p* < 0.05, ** *p* < 0.01, Student’s *t*-test. Bars = 100 μm (A) and 50 μm (B and D).

Finally, the differentiation of excitatory neurons was assessed by studying the layer-specific markers Ctip2 (layer V) and Mef2c (layers II-IV). *Dyrk1a^+/-^* mutants have fewer Ctip2^+^ neurons and more Mef2c^+^ neurons than their WT littermates (Fig. 3B and C). As the distinct types of neocortical excitatory neurons display different electrophysiological properties and project to distinct target areas (Molyneaux et al., 2007), the alteration in the proportions of these neurons could modify the final wiring of the brain in the *Dyrk1a^+/-^* mutants.

### 3.4. Alterations to the synaptic circuitry in the neocortex of adult*Dyrk1a*+/- mice

Around 10-15% of neocortical neurons are interneurons that make local connections and use γ-aminobutyric acid (GABA) as a neurotransmitter (Markram et al., 2004). These neurons maintain the stability of cortical networks and they modulate network activity through synaptic inhibition (Somogyi et al., 1998). In fact, disturbances to the excitation/inhibition ratio (E/I imbalance) in relevant brain circuits have been proposed as a common pathogenic mechanism in ASD (Rubenstein, 2010).

We have previously shown that the brain of adult *Dyrk1a^+/-^* mice present altered levels of proteins related to the glutamatergic and GABAergic systems (Souchet et al., 2014). Based on these data, we wondered whether the defects in cortical development displayed by the *Dyrk1a^+/-^* mice results in alterations in the adult circuitry. To this aim, we analyzed the excitatory (NeuN^+^/GABA^-^) and inhibitory (GABA^+^) neurons in the SSC of adult animals (Fig. 4A and B). The number of excitatory neurons in *Dyrk1a^+/-^* mutants increased in both the internal and external neocortical layers. The heterozygous mice also had more GABAergic neurons than their WT littermates, although the difference between these genotypes was only significant in the internal layers (Fig. 4B). The density of excitatory and inhibitory synapses was estimated in adult neocortices using antibodies against presynaptic markers VgluT1 (vesicular glutamate transporter 1) or VGAT (vesicular GABA transporter), and the postsynaptic markers Homer or Gephyrin. There was a significant increase in excitatory synapses in *Dyrk1a^+/-^* mutants in the two layers examined (IV and VI), while the density of inhibitory synapses in the mutant mice was similar to that in WT animals (Fig. 4C and D). The results suggest that the E/I imbalance in *Dyrk1a^+/-^* mice may provoke over-excitation in the cerebral cortex and related brain areas, contributing to the epileptic activity and ASD-related behavior in these animals.

**Fig. 4.**
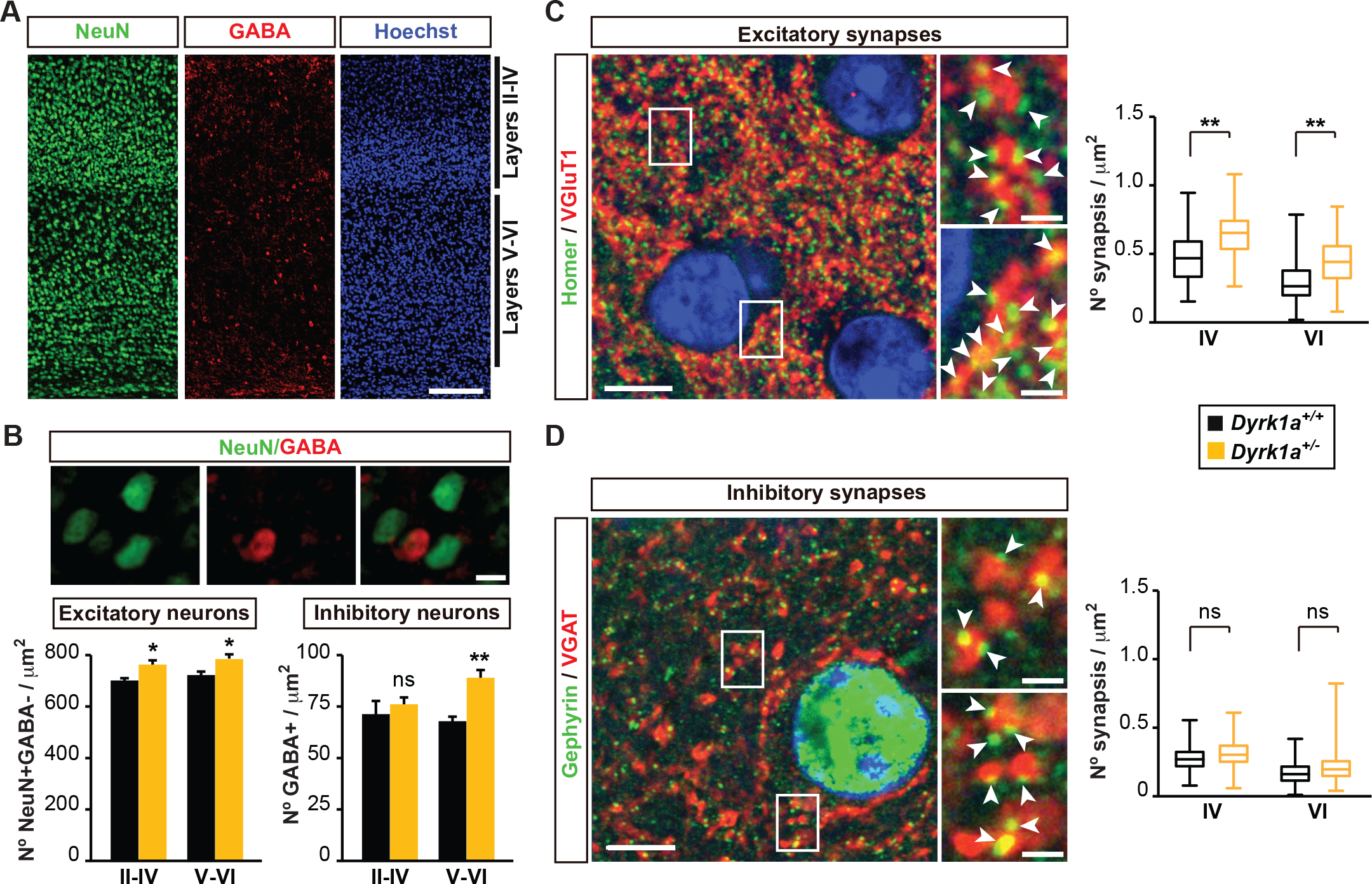
Adult *Dyrk1a*^+/-^ mice have a higher density of cortical neurons and synapses. (A) Representative images of the SSC from a *Dyrk1a*^+/+^ mouse section immunolabeled for NeuN and GABA, and counterstained with Hoechst nuclear dye. (B) Images showing NeuN^+^ and GABA^+^ neurons in the SSC, and histograms with the quantification of NeuN^+^/GABA^-^ (excitatory) and GABA^+^ (inhibitory) neurons in the external (layers II-IV) and internal (layers V-VI) layers of a 350 μm wide column of the SSC (n=4 animals each genotype; ns= not significant, \**p* < 0.05, \*\**p* < 0.01, Student’s *t*-test). (C, D) Representative confocal images of the SSC from a *Dyrk1a*^+/+^ mouse immunolabeled for VGluT1 and Homer (C), or for VGAT and Gephyrin (D), with the nuclei labeled with DAPI (blue). White rectangles indicate the area magnified in the images on the right. The white arrowheads point to interactions between presynaptic and postsynaptic markers. Box plots show the synapse densities between the first and third quartiles, the line in the boxes corresponding to the median synapse density (see Supplementary Fig. 4 for the automatic quantification of synapses). Excitatory synapses, layer IV (*Dyrk1a*^+/+^, n=639; *Dyrk1a*^+/-^, n=661) and layer VI (*Dyrk1a*^+/+^, n=639; *Dyrk1a*^+/-^, n=643). Inhibitory synapses, layer IV (*Dyrk1a*^+/+^, n=601; *Dyrk1a*^+/-^, n=539) and layer VI (*Dyrk1a*^+/+^, n=616; *Dyrk1a*^+/-^, n=535) (6 animals per genotype; \*\**p* < 0.01, ns=not significant, nested one-way ANOVA). Bars = 200 μm (A), 10 μm (B), and 5 μm and 1 μm (left and right photographs respectively in C, D).

### 3.5. Transcriptome alterations in the postnatal *Dyrk1a*^+/-^ cerebral cortex

The amount of Dyrk1a protein in the mouse developing neocortex increases during neurogenesis, reaching maximum levels in late embryonic development and during the first postnatal week (Dang et al., 2018). At these developmental stages, neural precursors shift their status and become gliogenic (Kriegstein and Alvarez-Buylla, 2009), while neurons extend neurites and synaptogenesis begins (Li et al., 2010). These processes are coordinated by transcriptional programs that are dynamically regulated during development (Telley et al., 2016), and it is possible that alterations to these programs might contribute to the early neurological phenotype in *Dyrk1a* haploinsufficiency. To test this possibility, we compared the transcriptional profile of the cerebral cortex in *Dyrk1a*^+/+^ and *Dyrk1a*^+/-^ mice at P0 and P7 (Supplementary Fig. 7A for the microarray experimental design and Fig. 5A for the comparisons performed). The comparison of the Dyrk1a expression profile during cortical development in *Dyrk1a*^+/+^ and *Dyrk1a*^+/-^ mice confirmed that *Dyrk1a* expression was reduced by half in the cortices of *Dyrk1a*^+/-^ mice at any time-point, reflecting *Dyrk1a* haploinsufficiency (Supplementary Fig. 6).

**Fig. 5.**
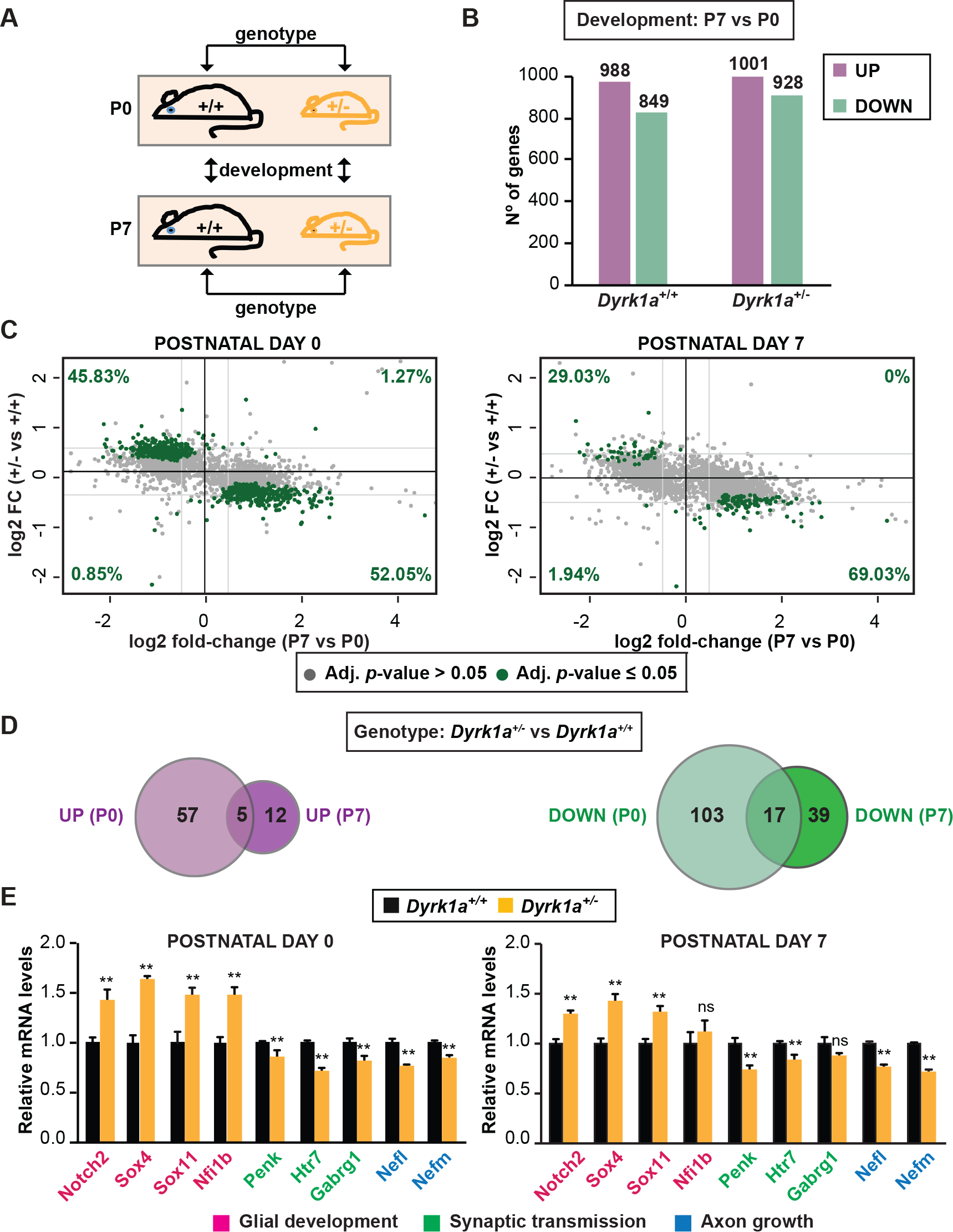
Transcriptome alterations in the neocortex of postnatal *Dyrk1a*^+/-^ mice. (A) Schematic representation of the comparisons performed to identify differentially expressed genes during development or between genotypes. (B) Transcriptional changes in the P7 and P0 cortex from *Dyrk1a*^+/+^ or *Dyrk1a*^+/-^ mice (adj. *p* ≤ 0.05; -1.4 ≤ FC ≥ 1.4). (C) Dot plots represent the changes in gene expression between genotypes (+/- *vs* +/+) at P0 and P7, and between the developmental stages (P7 *vs* P0) in the WT samples (+/+). Each dot represents a gene probe: green dots indicate probes with significant changes between genotypes (adjusted *p*-value <0.05); grey dots genes with no significant change. The numbers in green correspond to the percentage of green dots in each quadrant. Light grey lines indicate the log2 fold-change equal to ± 0.4. (D) The Venn diagrams show the genes up-regulated or down-regulated in comparisons of *Dyrk1a*^+/-^ with *Dyrk1a*^+/+^ mice at the two developmental time points studied, with the overlap indicating genes altered at both P0 and P7 (adj. *p* ≤ 0.05; -1.4 ≤ FC ≥ 1.4). (E) Validation by RT-qPCR of selected genes with altered expression in *Dyrk1a*^+/-^ animals. The graphs represent the mRNA expression relative to that in WT animals arbitrarily set as 1. RT-qPCR was used for *Nfi1b*, *Penk*, *Htr7*, *Nefl* and *Nefm* (n=4-9 animals each condition), and LDA for *Gabrg1, Notch2*, *Sox4* and *Sox11* (n=4 animals each condition; ns=not significant, \**p* < 0.05, Student’s *t*-test).

The heatmap representation of the microarray expression data showed that the developmental effect on gene expression was more relevant than the effect of the reduction in *Dyrk1a* dose (Supplementary Fig. 7B). In fact, the analysis of genes differentially expressed between P7 and P0 in *Dyrk1a*^+/-^ cortex showed numbers similar to the WT samples (Fig. 5B; see also Supplementary Fig. 8 for validation of the microarray results and Supplementary Dataset 1). A large proportion of the genes that were differentially expressed during development in the WT samples changed in the same direction in *Dyrk1a* mutant samples (Supplementary Fig. 7B). However, the expression of many of the genes that were up-regulated during development in the WT samples was diminished in *Dyrk1a^+/-^* samples and *vice versa* (Fig. 5C). These results suggest that *Dyrk1a* haploinsufficiency does not have a strong impact on the transcriptional program that drives postnatal cortical development but rather, that it might affect the time-course of the repression/activation of transcription. Nevertheless, the expression of a subset of genes was significantly altered between the two genotypes at P0 (182 genes) or P7 (73 genes), with changes greater or less than 1.4 fold (Fig. 5D).

Pathway enrichment analysis of the genes differentially expressed when comparing P0 and P7 WT cortices showed that the up-regulated genes at P7 were associated with synaptic transmission, while the down-regulated ones were enriched in cell cycle-associated functions (Supplementary Fig. 9). These data are consistent with the developmental changes occurring in the neocortex, since differentiating neurons acquire terminal features. Indeed, at these stages most pluripotent progenitors in the ventricular zone lose their capacity to self-renew or become less proliferative, producing cells of the glial lineage (Kriegstein and Alvarez-Buylla, 2009). When analyzing the genes with transcriptional changes in the *Dyrk1a*^+/-^ samples compared to the WT ones, no particular enrichment was associated with the up-regulated genes in the *Dyrk1a*^+/-^ animals, although known regulators of gliogenesis were identified in this set, such as *Sox4* or *Sox11* (Fig. 5E), suggesting a possible defect in the onset of glial cell production in the *Dyrk1a*^+/-^ mutant model. By contrast, the down-regulated set was enriched in genes whose products are located in dendrites (*Calb1*, *Penk*, *Thy1*) and/or that are involved in axon growth (*Nefl*, *Nefm*) or synaptic transmission (*Gabra*, *Gabrg1*, *Htr*: Fig. 5E and Supplementary Dataset 2). Hence, the transcriptional alterations detected further support that neuron activity is altered in the neocortex of *Dyrk1a*^+/-^ mutant pups.

## 4. Discussion

In this study, we analyzed a series of *DYRK1A* missense mutations, most of which are *bona-fide* LoF mutants that render the DYRK1A proteins enzymatically inactive. Moreover, the positive correlation between activity and protein accumulation might indicate that the presence of kinase-dead proteins has an added harmful effect. Very recently, the analysis of 5 DYRK1A missense mutants within the catalytic domain (L245R, D287V, L295F, F308V and S311F) has been published (Widowati et al., 2018). Our results concur and further expand this analysis by including all missense mutations in the catalytic domain of DYRK1A published to date. Some of these mutations lie in regions predicted to be essential, as the case of K188 in the ATP-binding domain or D287 and F308 (HCD- and DFG-motifs, respectively), which are involved in the catalytic reaction. There is a particular accumulation of LoF mutations within the P+1 loop and the α-helix F. In addition, though the number of missense mutations is still low, some residues can be considered as hot spots either because the mutation has been reported in two different patients (S346P: Bronicki et al., 2015; Deciphering Developmental Disorders, 2015; R467Q: (Evers et al., 2017; Posey et al., 2016)) and/or because the same residue is mutated to different amino acids (D287V and D287Y: Deciphering Developmental Disorders, 2015; Zhang et al., 2015; S346P and S346F: Bronicki et al., 2015 and ClinVar SCV000712522).

Somewhat unexpectedly, the enzymatic activity of several missense mutants was indistinguishable from that of the WT kinase and in some cases, even an increment in activity was detected. We cannot exclude that the activity of some of these mutants could be the reflection of an altered substrate choice; alternatively, they might lead to other alterations in DYRK1A function affecting its final biological activity. A third possibility could be that they represent pathogenic variants of DYRK1A hyperactivity, based on the links of DYRK1A overexpression with certain DS pathological traits (Becker et al., 2014). Of note, the hyperactive G486D mutation was identified in a patient with macrocephaly (Dang et al., 2018), in contrast with the DHS patients with LoF mutations in which microcephaly is, generally, observed (Supplementary Table 1). However, the results might also mean that these mutations do not provoke any negative effects. Indeed, the effects of the catalytic missense A195T and L259F mutants have been evaluated in cultured cortical neurons, where their impact on neurite extension was similar to that of the WT neurons (Dang et al., 2018). Therefore, and for the catalytically active mutants, there is the possibility that the genetic variant in the *DYRK1A* gene is a benign variant and therefore not responsible for the clinical phenotype.

In ASD and ID syndromes, alterations to dendrite morphogenesis and synapse formation are common (de la Torre-Ubieta et al., 2016). There is evidence that DYRK1A regulates the actin cytoskeleton and microtubule dynamics, thereby contributing to the development and maintenance of neurites and dendritic spines (Martinez de Lagran et al., 2012; Ori-McKenney et al., 2016; Park et al., 2012). Indeed, the morphology of the dendritic arbor in neurons of the motor cortex is altered in *Dyrk1a*^+/-^ mice and in transgenic mice overexpressing *Dyrk1a* (Benavides-Piccione et al., 2005; Martinez de Lagran et al., 2012). These, together with the detrimental effect of *DYRK1A* truncated mutations in neuronal dendritic and spine growth (Dang et al., 2018) suggested that postnatal neocortical development is relevant to the neuropathology of DHS. However, our results indicate that *Dyrk1a* haploinsufficiency also affects neuron production in the developing neocortex leading to alterations to the number and proportion of the excitatory neuron subtypes. Considering that each of the many distinct excitatory neural subtypes fulfills a particular function (Jabaudon, 2017), subtle alterations in the proportion of these neurons would influence the activity of the synaptic circuits formed during postnatal development. In this context, the different transcriptomic profiles of the postnatal *Dyrk1a*^+/-^ neocortex could be the consequence of its different neuronal composition. However, given that DYRK1A can directly or indirectly regulate transcription (Aranda et al., 2011; Di Vona et al., 2015), *Dyrk1a* haploinsufficiency might specifically modify the transcriptional programmes in differentiating neocortical neurons. In any case, the transcriptome analysis highlighted the weaker expression of genes involved in neuritogenesis and synaptic activity in the postnatal *Dyrk1a*^+/-^ neocortex. Therefore, both the reduction in gene expression and the direct activity of DYRK1A on proteins implicated on neuritogenesis should be considered as factors that contribute to the neurological alterations in *Dyrk1a*^+/-^ mutant mice.

Adult *Dyrk1a*^+/-^ mice experience epileptic activity and behavior deficits similar to those observed in other ASD mouse models in which excitatory and/or inhibitory synaptic circuits are perturbed (Lee et al., 2017). In fact, altered production and/or migration of excitatory and/or inhibitory cortical neurons affects the E/I balance in mature circuits, usually making them epileptogenic (Bozzi et al., 2012; Rubenstein, 2010). *Dyrk1a*^+/-^ mutants exhibit an excess of excitatory neurons and synapses in the neocortex, and these animals also show an excess of inhibitory neurons in the internal neocortical layers. Thus, we propose that both excitatory and inhibitory neurons contribute to the E/I imbalance underlying the epileptic activity, ASD-like behavior and cognitive deficits associated with DHS.

## Abbreviations

ASD, autism spectrum disorder; BrdU, bromodeoxyuridine; DHS, *DYRK1A* haploinsufficiency syndrome; DS, Down syndrome; DYRK, dual-specificity tyrosine-regulated kinase; EEG, electroencephalogram; E/I, excitation/inhibition; GABA, λ-aminobutyric acid; GFP, green fluorescent protein; ID, intellectual disability; IVK, *in vitro* kinase; LDA, low-density array; LoF, loss-of function; SSC, somatosensorial cortex; UsV, ultrasonic vocalization; WT, wild-type.

## Acknowledgements

The microarray studies were performed at the Genomics Platform of the Institut de Recerca Vall d’Hebron (VHIR). We thank Fatima Nuñez (VHIR’s Genomics Platform) for her help in designing the microarray experiment, Alex Sánchez and Ricardo González (VHIR’s Statistics and Bioinformatics) for the statistical analysis of the microarray data, Alexis Rafols (CRG Histology Unit) and Carmen Badosa for technical support, and Mark Sefton for English editorial work.

## Supplementary Materials and Methods

### Western blots

Samples were resolved by SDS-PAGE and the proteins were transferred onto nitrocellulose membranes (Amersham Protran). The membranes were blocked for 30 min at room temperature with 10% skimmed milk (Sigma) diluted in 10 mM Tris-HCl [pH 7.5], 100 mM NaCl (TBS) plus 0.1% Tween-20 (TBS-T), and then incubated overnight at 4°C with the primary antibodies (Supplementary Table 3) diluted in 5% skimmed milk in TBS-T. After several washes, the membranes were incubated for 45 min at room temperature with horseradish peroxidase-conjugated secondary antibodies (DaKo) diluted in 5% skimmed milk in TBS-T. Antibody binding was detected by enhanced chemiluminiscence (Western Lightning Plus ECL, Perkin Elmer), which was analyzed on a LAS-3000 image analyzer (Fuji PhotoFilm). The antibody signal was quantified with the ImageQuant software. For relative protein accumulation, total cell extracts were used by resuspending the cell pellets in SDS loading buffer. For quantification, the signal of the HA-antibody was corrected with that of the GFP-antibody in co-transfections with pEGFP-C1 (Fig. 1E).

### Microarray experiments

The experimental set up is depicted in Supplementary Fig. 6A. Total RNA was prepared from the cerebral cortex of P0 and P7 mice, obtained from two litters (*Dyrk1a^+/+^*, n=4; *Dyrk1a*^+/-^, n=4). Double stranded cDNA was synthesized from 100 ng RNA using the Two Cycle cDNA Synthesis Kit (Affymetrix), according to the manufacturer’s instructions. After cRNA purification and fragmentation, the cRNA quality was checked (3’/5’ ratio of probe sets <1.5, for *Gapdh* and *Actb* probes). The fragmented cRNA (15 μg) was added to the hybridization mix and used to fill the Affymetrix Mouse GeneChip 430 2.0 Array cartridge (over 39,000 mouse genes). The arrays were hybridized at 60°C for 16 h with rotation in the Affymetrix GeneChip Hyb Oven 640, washed and marked with streptavidin R-phycoerythrin following the Affymetrix protocol. After washing, the arrays were scanned in an Agilent G3000 GeneArray Scanner and the images were processed with Microarray Analysis Suite 5.0 (Affymetrix). Raw values obtained from the .CEL files were pre-processed using the Robust Multiarray Averaging Method (Irizarry et al., 2003), and the resulting normalized values were used for all subsequent analyses. The data were first subjected to non-specific filtering to remove low signal genes (those whose expression was below the third quartile of all expression values) and low variability genes (those whose standard deviation of the expression was below the third quartile of all standard deviations). All the statistical analyses were performed with the “R” statistical software and the libraries developed for microarray data analysis using the Bioconductor Project (www.bioconductor.org). Probes with an adjusted *p*-value ≤ 0.05 that mapped to a known mouse gene were selected for the results shown in Fig. 5 (Supplementary Dataset 1).

### RT-qPCR

The cDNA was synthesized from 1 μg of total RNA using Superscript II retrotranscriptase (Invitrogen) and random hexamers. Real-time qPCR was carried out on the Lightcycler 480 platform (Roche), using the SYBR Green I Master kit (Roche) and cDNA diluted 1:10, mixing the reagents according to the manufacturer’s recommendations. In general, the PCR conditions were: one cycle 95°C 5 min; 45 cycles 95°C 10 sec, 60°C 10 s and 72°C 10 s; one cycle 95°C 5 sec and 65°C 1 min. Primers for RT-qPCR, were designed using the “Primer3” tool (frodo.wi.mit.edu/cgi-bin/primer3/primer3) to obtain 80-200 bp amplicons and optimal annealing temperatures between 59 and 61°C for each primer pair (Supplementary Table 4). We used *Ppia* as a reference gene for data normalization. Each sample was assayed in triplicate and the Ct (threshold cycle) was calculated using the relative quantification of the second derivative maximum method with the Lightcycler 480 1.2 software (Roche).

For qPCR experiments on a Low Density array (LDA, Applied Biosystem), cDNA was synthesized from 0.5 μg of total RNA using MultiScribe retrotranscriptase and using random primers (High Capacity cDNA Reverse Transcription Kit, Applied Biosystem), according to the manufacturer’s recommendations. For quantitative analysis, the cDNAs were diluted 1:10 in H_2_O, added to TaqMan^®^ Universal PCR Master Mix (Applied Biosystem) in a 1:1 ratio and loaded onto the LDA, pre-loaded with the selected TaqMan^®^ Gene Expression Assays (Supplementary Table 5). The TaqMan Array was run on the 7900HT system (Applied Biosystem) following the manufacturer’s instructions, and the data were analyzed with the SDS software (Applied Biosystem), based on the Pfaffl Method (Pfaffl, 2001). We used *Ppia* and *Actb* as the reference genes for data normalization.

### Computational analysis

The differentially expressed genes were selected based on a linear model analysis with empirical Bayes modification for the variance estimates (Smyth, 2004) and the p-values were adjusted using the Benjamini-Hochberg method (Benjamini and Hochberg, 1995). We used hierarchical clustering with Euclidean distances to form the groups of differentially expressed genes and the heat maps to visualize them. The annotation of the Affymetrix dataset was done with bioDBnet (biodbnet-abcc.ncifcrf.gov; Mudunuri et al., 2009). The analysis of transcription factor and pathway enrichment was performed using the web server Enrichr (amp.pharm.mssm.edu/Enrichr; Kuleshov et al., 2016) and its associated visualization tools (Tan et al., 2013). The Ensemble (www.ensembl.org) and NCBI (www.ncbi.nlm.nih.gov) public databases were used to integrate the information regarding gene sequences and functions.

### Behavioral Testing

#### Social interactions

Social interactions between two unfamiliar mice were evaluated using a modification of the reciprocal social interaction test (Lin and Hsueh, 2014). In brief, the subject animal was placed in a standard housed cage (12.5 cm x 17.5 cm x 32.5 cm) with clean bedding and after 5 min of habituation, a male of a different litter but on the same genetic background and age was introduced into the cage. The interactions between the two mice were videotaped during 5 min and the same intruder mouse was used in all tests. We scored the time spent in the social interactions (nose-to-nose sniffing and nose-to-anus sniffing) that were initiated by the subject mouse and the time spent by this mouse performing stereotyped repetitive behaviors (digging and grooming).

#### Marble-burying

The marble-burying test was performed as described previously (Thomas et al., 2009). In brief, mice were placed in a cage (17.5 cm x 24.5 cm x 42.5 cm) that had 12 black glass marbles (15 mm diameter) evenly spaced in a 2 x 6 grid on top of 5 cm of clean bedding. The mice were videotaped for 20 min, and we scored the number of marbles buried (90% or more covered) and the time spent by the mouse performing stereotyped repetitive behaviors (digging and grooming).

#### Ultrasonic vocalization (UsV)

UsV was measured in male and female pups at P3, P6, P9 and P12 (4 litters, 17-11 animals each genotype). The dam was removed from a temperature-controlled home cage where the pups remained. The pups were then removed individually from the cage and transported in a dish into a hood equipped with a UsV recorder (Avisoft) where they were recorded for 5 min. Room temperature was maintained at 21°C and the body temperature was measured with an axillary probe after the 5 min test. The data was analyzed with Avisoft SASLab Pro software.

### Immunostaining

To prepare cryotome sections, brains were cryoprotected with 30% sucrose in phosphate-buffered saline (PBS), while for vibratome sections brains were embedded in 4% agarose in H_2_O. For immunohistochemistry, cryotome sections (14 μm) were collected on Starfrost precoated slides (Knittel Glasser) and distributed serially. For immunofluorescence, free-floating cryotome and vibratome sections (30-45 μm) were distributed serially in 48-well plates. For bromodeoxyuridine (BrdU) antibodies the sections were treated for 30 min with 2N HCl at 37°C before blocking (Najas et al., 2015). Immunohistochemistry was performed following the avidin-biotin-peroxidase method (Vectastain ABC kit, Vector Labs, Burlingame, CA, USA) as described previously (Barallobre et al., 2014). In this case, samples were stained with Nissl (0.1% Cresyl violet acetate) to visualized non-immunopositive cells. For immunofluorescence, the sections were blocked for 1 h at room temperature in PBS containing 0.2% Triton-X100 and 10% fetal bovine serum (FBS), and probed for 18 to 96 h at 4°C with the primary antibodies (Supplementary Table 6) diluted in antibody buffer (PBS with 0.2% Triton-X100 and 5% FBS). For some antibodies, an antigen retrieval treatment was performed before blocking. The sections were then washed and the primary antibodies were detected using Alexa-555 or Alexa-488 conjugated secondary antibodies (1:1000; Life Technologies). Cell nuclei were stained with Hoechst or 4′,6-diamidino-2-phenylindole,(DAPI; Sigma-Aldrich). For antibodies against Gephyrin and Homer, signal amplification was required and thus, after washing the sections they were incubated with the corresponding biotinylated secondary antibody (1:200; Vector Labs), and then with Alexa-488 conjugated streptavidin (Life Technologies). In all cases, the specificity of the immunoreaction was assessed by omitting the primary antibody.

### Cell and synapse counts

Wide-field and confocal microscope images were taken from the somatosensorial cortex (SSC) and processed using Image J/Fiji (Rasband, W.S., National Institutes of Health, Bethesda, Maryland, USA; imagej.nih.gov/ij).

#### Cell counts

Cells in postnatal animals were counted in 150 μm wide radial columns (NeuN^+^ cells), in 350 μm wide radial columns (Citp2^+^ cells and Mef2c^+^ cells) or in 500 μm wide radial columns (BrdU^+^ cells) of the SSC. Labeled cells in adult animals were counted in 150 μm wide radial columns (NeuN^+^/GABA^-^ cells) or in 350 μm wide radial columns (GABA^+^ cells) of the SSC. All cell counts were obtained blind from images at the same rostro-caudal level of at least 3 sections per animal: from the beginning of the hippocampal commissure to the beginning of the hippocampus in P7 mice, and from bregma -0.22 to bregma -0.94 (Paxinos and Franklin, 2001) in adult mice.

#### Synapse counts

Tissue samples used for synapse counts where mounted in 2,2’-thiodiethanol adjusted to 1.51 refraction index in order to minimize chromatic shift. Synapses were counted in images from 3 sections at the same rostral-caudal level in each mouse (sections from bregma -0.22 to bregma -0.94). Twenty images (26 μm^2^) per hemisphere and layer were analyzed randomly from the SSC layers IV and VI on a Zeiss LSM780 confocal microscope equipped with a Plan Apochromat 63x oil (NA=1.4) objective. 2D confocal sections were acquired by sequential line scanning using the excitation lines 405, 488 and 561 and the respective detection bandwidths 415-491, 490-553 and 568-735. The scanning speed was set at 3.15 μs/pixel and the resolution was adjusted to 50 nm/pixel. All images were acquired under the same parameters to guarantee the signals captured remained within the limits of the dynamic range of detection. A specific algorithm was developed in macro language to automatically obtain the density of the synapses in all the images (see details in Supplementary Fig. 4).

---

**Supplementary Table 1.**
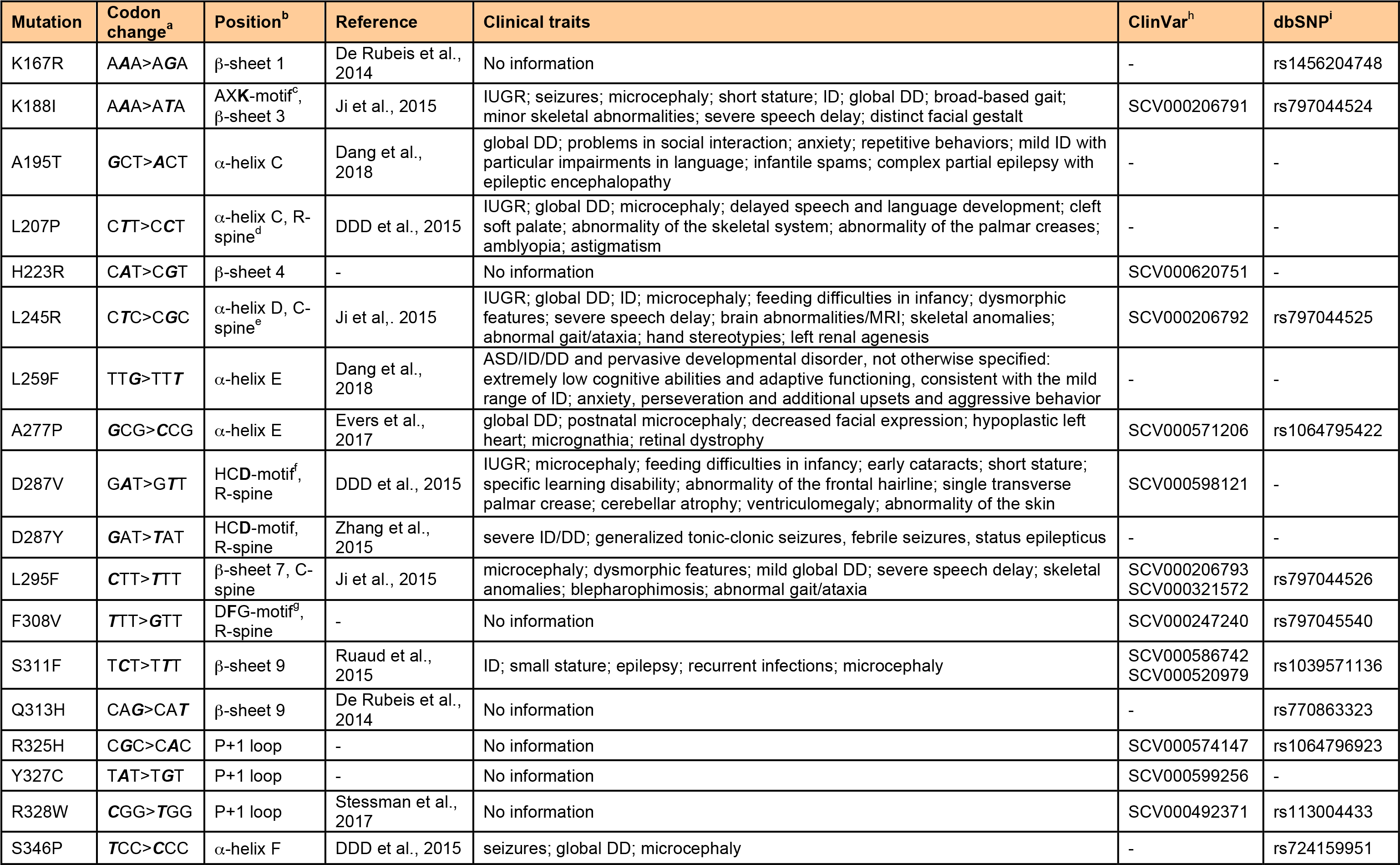
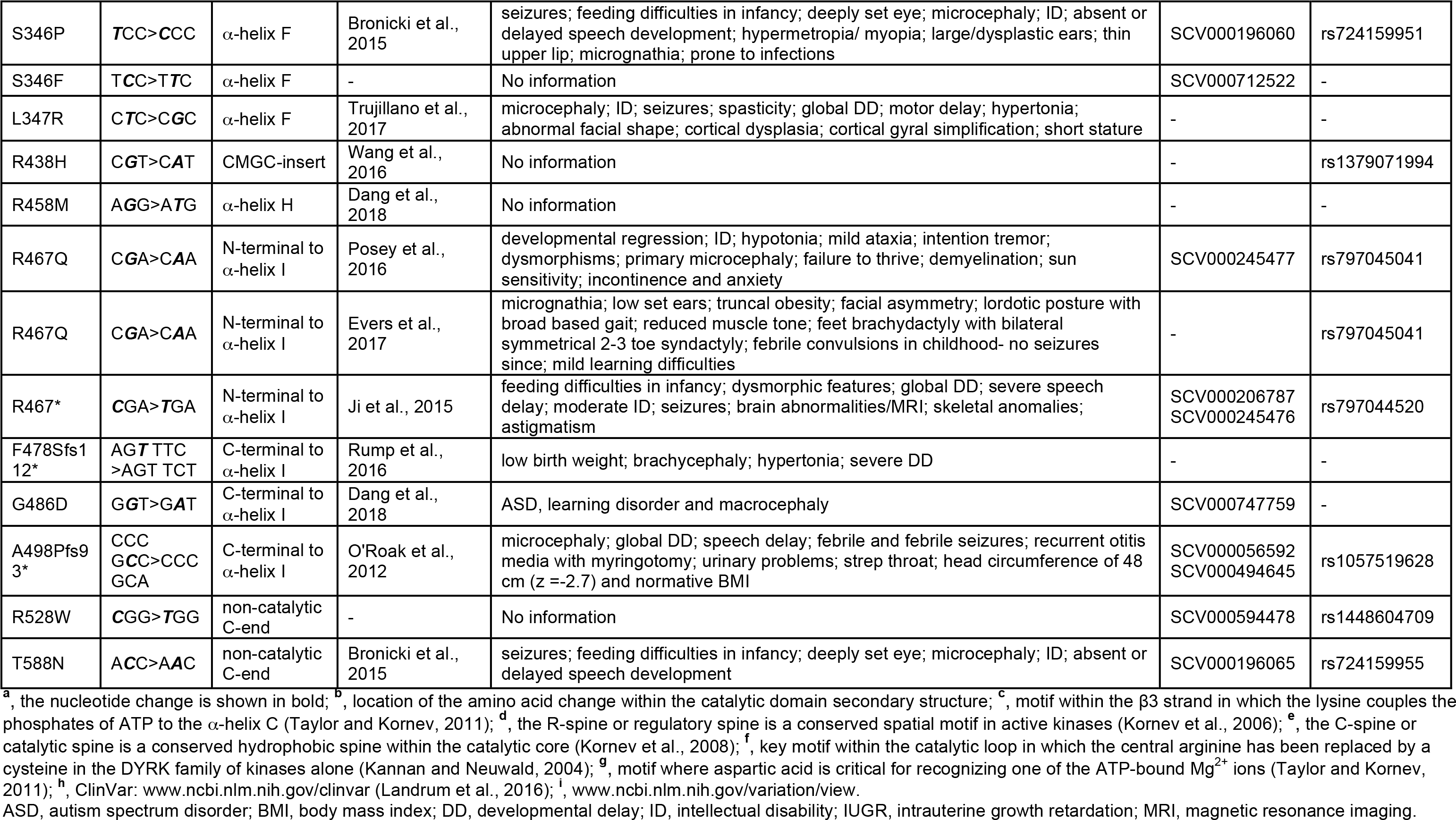
Mutations analyzed in this study.

**Supplementary Table 2.**
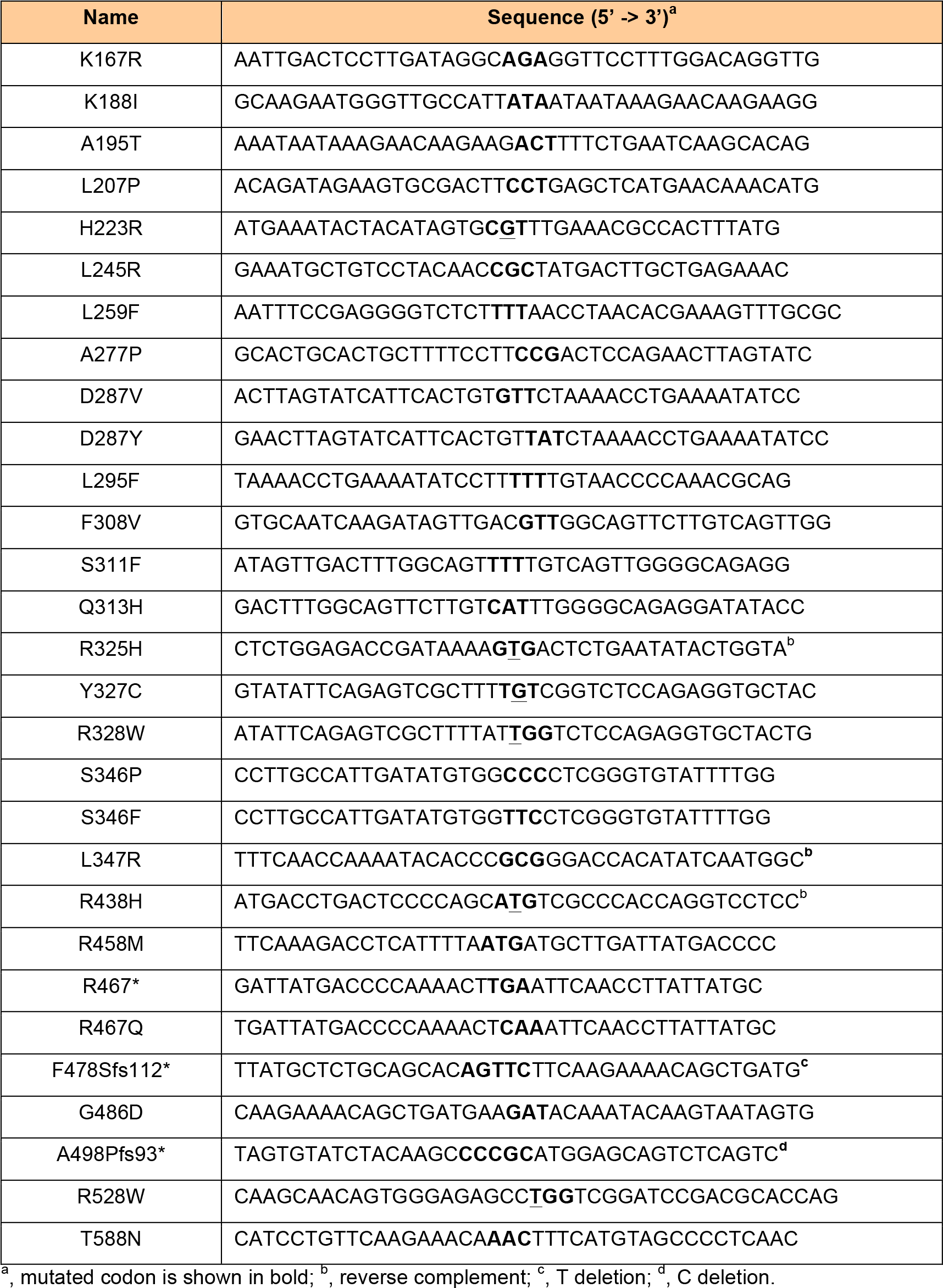
Primers used for site directed mutagenesis.

**Supplementary Table 3.** Primary antibodies used for immunoblotting.

| Antibody | Host | Source | Dilution |
| --- | --- | --- | --- |
| DYRK1A | Mouse | Abnova (H000001859) | 1/1000 |
| DYRK1A | Rabbit | Abcam (ab69811) | 1/1000 |
| GFP | Mouse | Clontech (632381) | 1/2000 |
| HA | Mouse | Covance (MMS-101R) | 1/2000 |
| phospho-Tyr (PY99) | Mouse | Santa Cruz (sc-7020) | 1/1000 |
| α-Tubulin | Mouse | Sigma (T6199) | 1/5000 |

**Supplementary Table 4.**
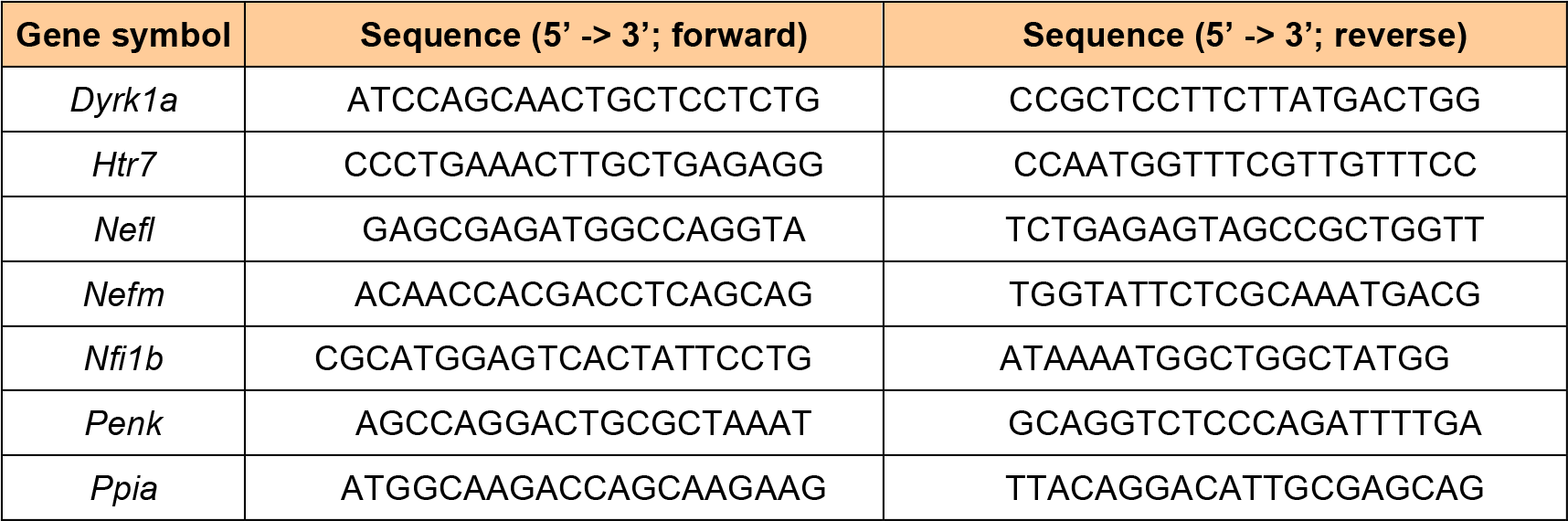
Primers used for qPCR.

**Supplementary Table 5.**
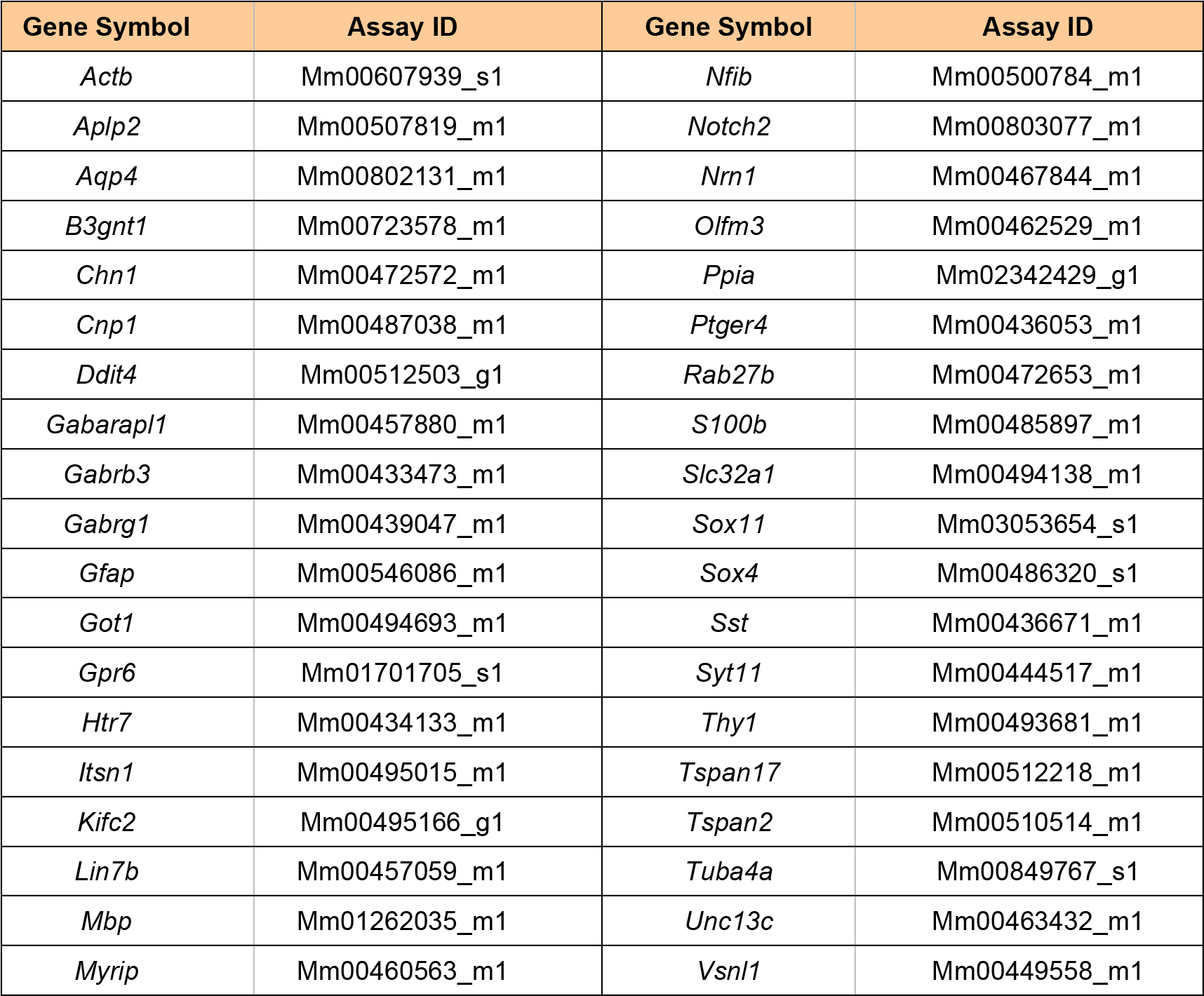
Probes used in the Low Density Array.

**Supplementary Table 6.** Primary antibodies used for immunostaining.

| Antibody | Host | Source | Dilution |
| --- | --- | --- | --- |
| BrdU | Rat | Serotec (OBT0030G) | 1:100 |
| Ctip2* | Rat | Abcam (ab18465) | 1:250 |
| GABA | Rabbit | Sigma-Aldrich (A2052) | 1:200 |
| Gephyrin* | Mouse | Synaptic Systems (147 011) | 1:500 |
| Homer* | Mouse | Thermo Scientific (MA1-045) | 1:1000 |
| Mef2c* | Rabbit | Abcam (ab64644) | 1:250 |
| NeuN | Mouse | Chemicon (MAB377) | 1:1000 |
| VGAT* | Rabbit | Synaptic Systems (131 003) | 1:400 |
| VGlut1* | Guinea-pig | Milipore (AB5905) | 1:700 |

\*Citrate pretreatment

## Supplementary Figure Legends

**Supplementary Fig. 1.**
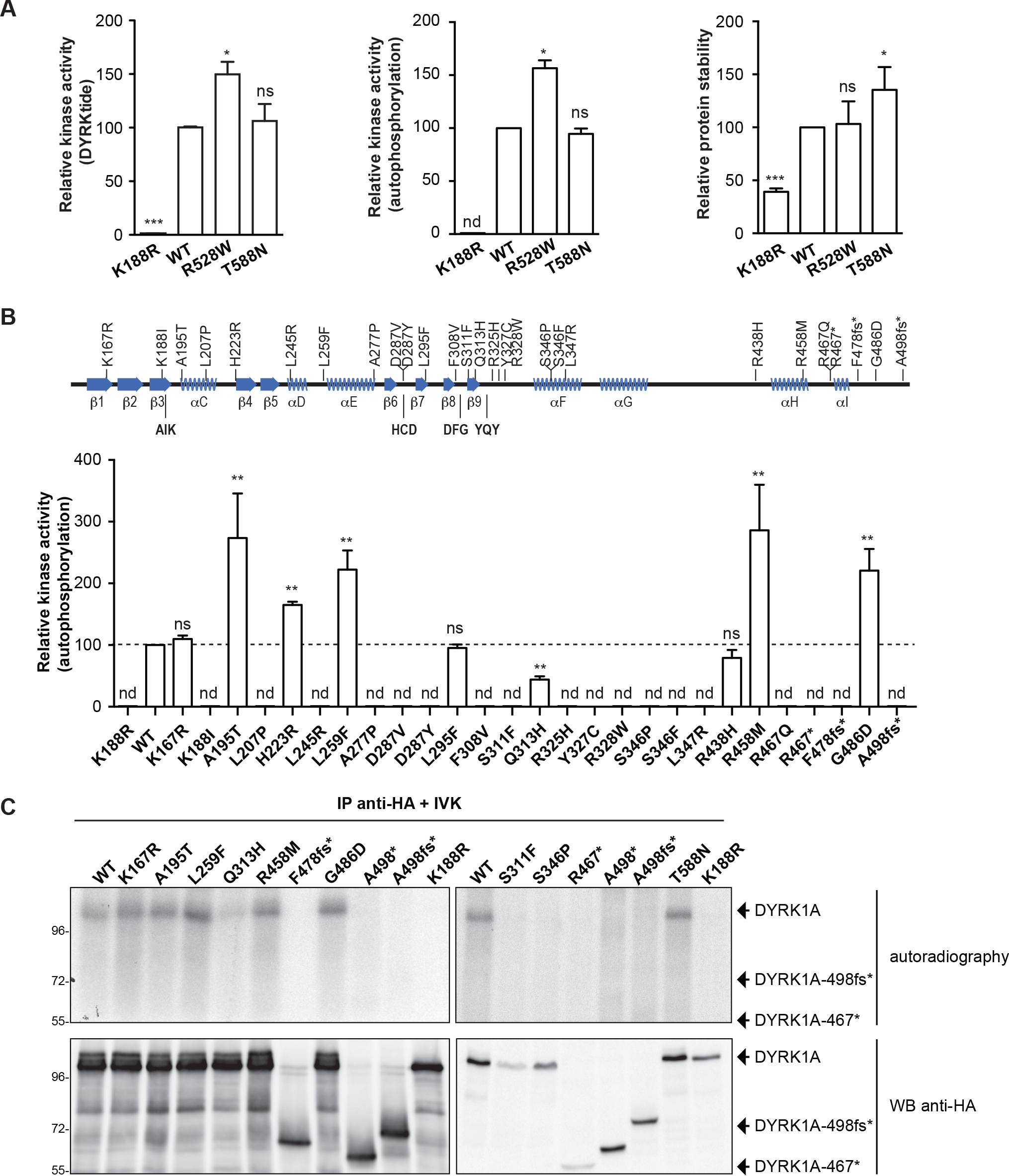
Biochemical analysis of *DYRK1A* mutants. (A) Analysis of the DYRK1A mutants outside the catalytic domain, R528W and T588N. The graphs show the mean ± SEM, with the WT values set arbitrarily as 100 (n=4, for DYRKtide phosphorylation; n=3, for autophosphorylation; n=3, for protein accumulation). (B) Analysis of the autophosphorylation activity of the DYRK1A mutants shown in Fig. 1. The graphs show the mean ± SEM, with the WT values set arbitrarily as 100 (n=3). In the authophosphorylation assays in A and B, nd refers to radioactive signals at the background level (ns= not significant, \**p* < 0.05, \*\**p* < 0.01, \*\*\**p* < 0.001, unpaired 2-tailed Mann-Whitney’s test). (C) Representative images of the autophosphorylation assays for selected DYRK1A mutants.

**Supplementary Fig. 2.**
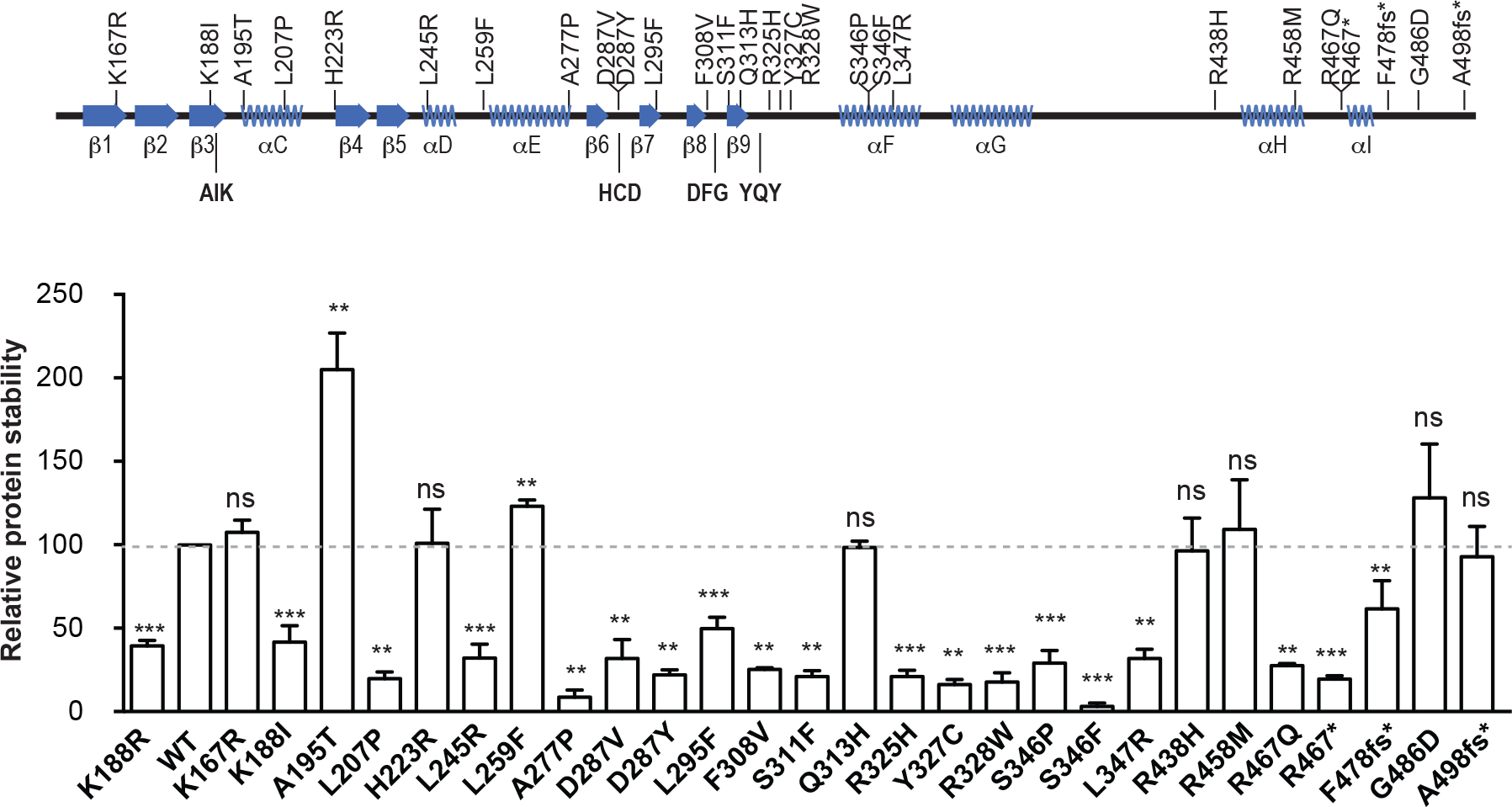
Protein accumulation analysis of *DYRK1A* mutants. Accumulation of DYRK1A mutants calculated as described in Fig. 1E, with the WT protein values set arbitrarily as 100 (mean ± SEM, n=3 independent experiments; ns= not significant, \*\**p* < 0.01, \*\*\**p* < 0.001, unpaired 2-tailed Mann-Whitney’s test).

**Supplementary Fig. 3.**
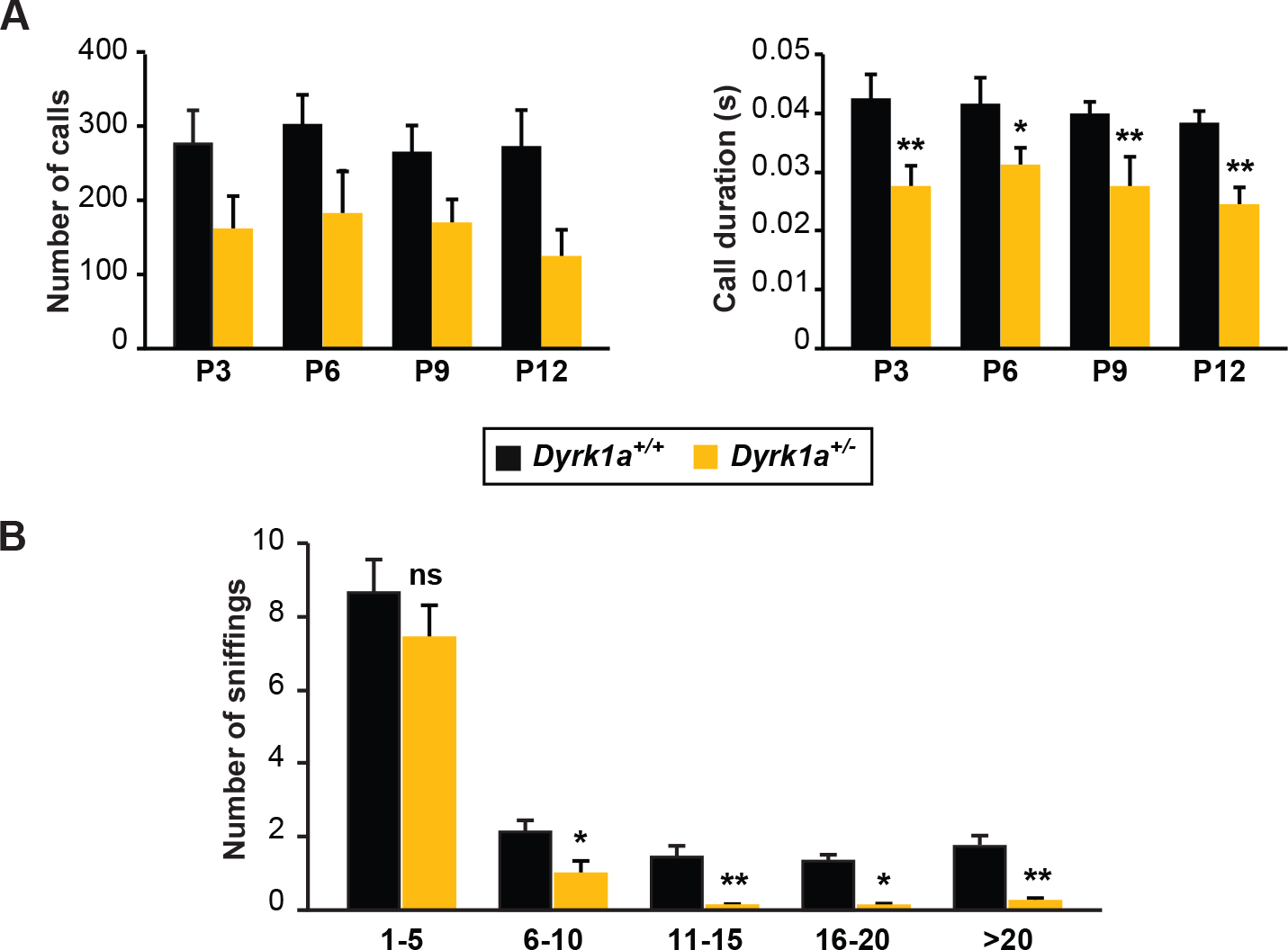
ASD-related phenotypes of *Dyrk1a*^+/-^ mice. (A) Number and duration of UsV calls recorded over 5 min from pups when separated from their mothers at the developmental stage indicated (*Dyrk1a^+/+^,* n=17; *Dyrk1a*^+/-^, n=11). UsV data was analyzed by two-way ANOVA followed by a Bonferroni post-hoc test. The differences between the genotypes were statistically significant: F_(1.94)_=13.25, p = 0.0004 (number of calls) and F_(1.92)_ =40.92, *p* < 0.0001 (call duration), two-way ANOVA. Differences in call duration were significant in the Bonferroni post-hoc test (\**p* < 0.05, \*\**p* < 0.01). (B) Social contacts (sniffing) performed by *Dyrk1a*^+/+^ and *Dyrk1a*^+/-^ mice in the social interaction test. The graph represents the number of sniffs (mean ± SEM) over 5 min distributed according to their duration (*X*-axis correspond to seconds). Note that *Dyrk1a*^+/-^ mice engaged in significantly fewer social contacts longer than 6 s: (n=12-15 mice each genotype; \**p* < 0.05, ** *p* < 0.01, Student’s *t*-test).

**Supplementary Fig. 4.**
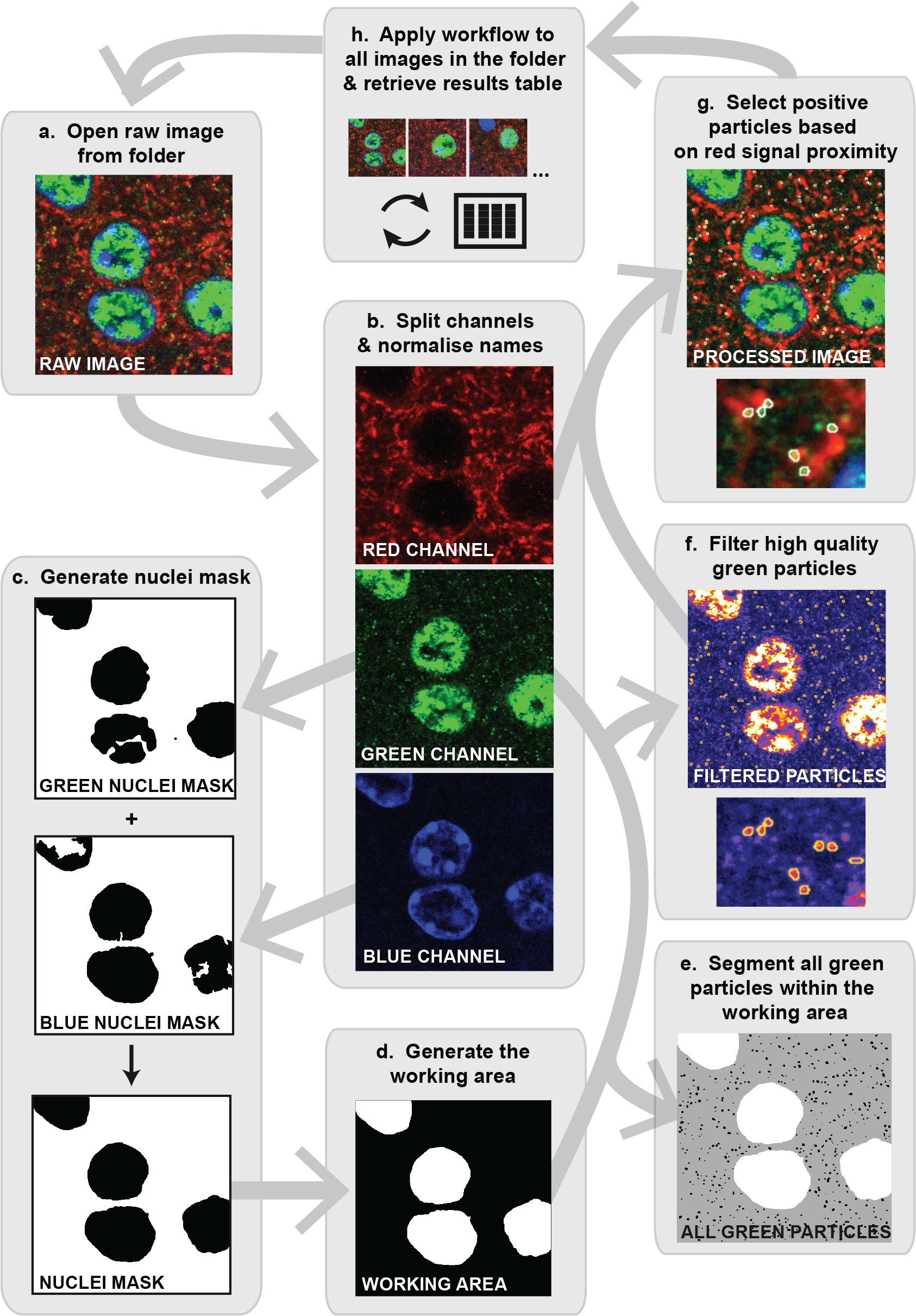
Scheme showing the workflow for the automatic quantification of synapses from confocal images using an image-processing algorithm written in Fiji macro language. (A) The macro opens a raw data image from a folder of choice (file with an extension *.lsm). (B) The algorithm splits the channels and normalizes their names to “Blue” (DAPI nuclear staining), “Green” (Alexa Fluor-488 for post-synaptic labeling of Homer or Gephyrin) and “Red” (Alexa Fluor-568 for pre-synaptic labeling of VgluT1 or VGAT). As Gephyrin (example in panel B) but not Homer is found in the nuclei, there were minor differences in the processing of the “Green” signal in some of the following steps. (C) The “Blue” channel is pre-processed to remove the noise by applying a Gaussian filter (Radius=2) and a Rolling Ball algorithm (Radius=80). An automatic threshold (Li) was then applied and the objects were filtered to generate a binary mask containing only the nuclei. In the case of Gephyrin^+^/VGAT^+^ images, the “Green” channel was pre-processed with a Gaussian filter (Radius=5), automatically thresholded (Huang) and the objects were filtered to generate a second mask containing only the nuclei. Both masks (Gephyrin and DAPI nuclear signals) were combined to obtain a single nuclear mask. (D) The inverted nuclear mask defines the working area, and this value was stored and used in the final calculation of synapse density, the selected area being used to estimate the median signal value of the “Red” channel. The median signal values for all the images in the data set were averaged to obtain a global median value, which was used at a later step as a reference to normalize the red channel. (E) To enhance spot discrimination, the original “Green” channel was filtered using the Laplacian of Gaussian (Radius=2) operator. After applying an automatic threshold (Otsu), the spots detected were filtered according to size (0.01-100 μm^2^) and circularity criteria (0.6-1), and used to produce a binary mask. Finally, the watershed algorithm was used to further separate contiguous particles. (F) The segmented post-synaptic particles in the original “Green” channel were further filtered according to the density of their intensity, and small and/or weak particles below a certain threshold were discriminated. (G) To count synapses, the “Red” channel was first translated to correct for the chromatic shift (x=0.87 and y=1.31: values calculated using fluorescent microspheres), it was pre-processed with a Median filter (Radius=2) and Rolling ball algorithm (Radius=120) to remove noise, and it was normalized by the global median signal calculated in D. The macro counts a synapse as every green particle selected that overlies at least one pixel with a relevant intensity in the red channel. A verification image is then generated that contains all three processed channels and the synapse events highlighted in white. (H) The macro analyses all the images included in a folder of choice and delivers a text file for each image that contains the image name, the median red intensity value, the working area value, the number of segmented green particles, the number of filtered green particles and the number of synapses. Synapse density was calculated by dividing the number of synapses by the value of the working area.

**Supplementary Fig. 5.**
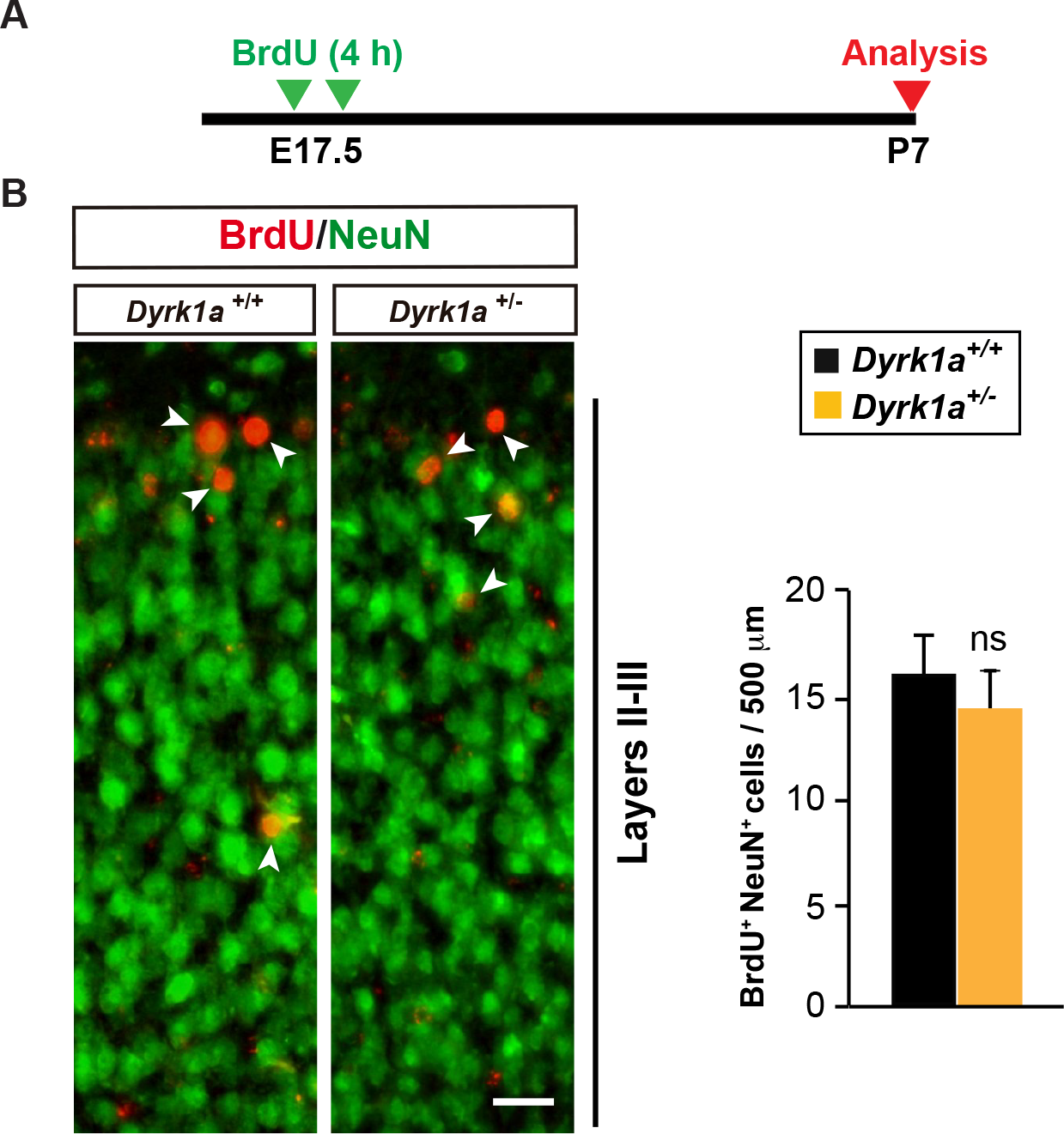
Neuron production in the dorsal telencephalon of E17.5 *Dyrk1a^+/+^* and *Dyrk1a*^+/-^ embryos. (A) Schedule of the BrdU-labeling protocol to estimate neuron production in the developing neocortex. (B) Representative images of the SSC of P7 *Dyrk1a^+/+^* and *Dyrk1a*^+/-^ animals showing BrdU^+^ cells immunopositive for the neuronal marker NeuN (arrowheads). Bar = 50 μm. The histogram represents the number of BrdU^+^/NeuN^+^ neurons in a 500 μm wide column of the neocortex (n=3 animals each genotype; ns=not significant, Student’s *t*-test). In both genotypes, the are few BrdU^+^/NeuN^+^ neurons and these neurons are positioned in the most external part of the neocortex.

**Supplementary Fig. 6.**
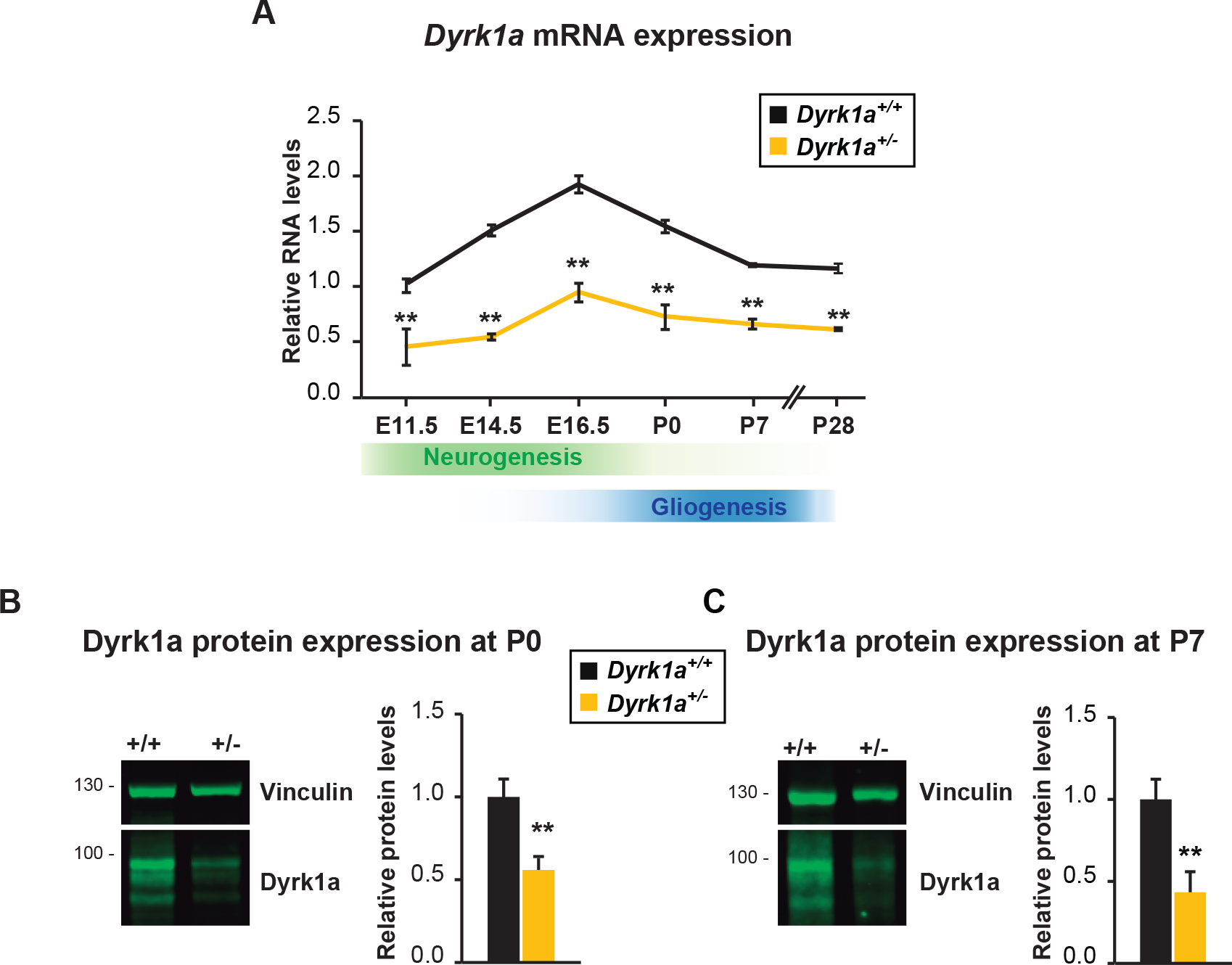
Dyrk1a expression in the developing cerebral cortex of *Dyrk1a*^+/-^ mutants and their WT littermates. (A) *Dyrk1a* mRNA in the developing cerebral cortex of *Dyrk1a^+/+^* and *Dyrk1a*^+/-^ embryos or postnatal animals. The graph represents the mRNA expression relative to the levels in E11.5 *Dyrk1a^+/+^* embryos set arbitrarily as 1 (n  3; \*\**p* < 0.01, Student’s *t*-test, +/- vs +/+ at any time point). Note that the reduced *Dyrk1a* expression in the mutants is maintained during embryonic and postnatal development. (B, C) Representative Western blots of extracts prepared from P0 (B) and P7 (C) cerebral cortices and probed with the antibodies indicated. The secondary antibody was detected by infrared fluorescence using the LI-COR Odyssey IR Imaging System V3.0 (LI-COR Biosciences, Cambridge, UK). The graphs represent the Dyrk1a protein normalized to Vinculin levels and expressed relative to the WT animals. Note that there is approx. 50% less Dyrk1a in *Dyrk1a*^+/-^ mice (n=3 mice each genotype): \*\**p* < 0.01, Student’s *t*-test.

**Supplementary Fig. 7.**
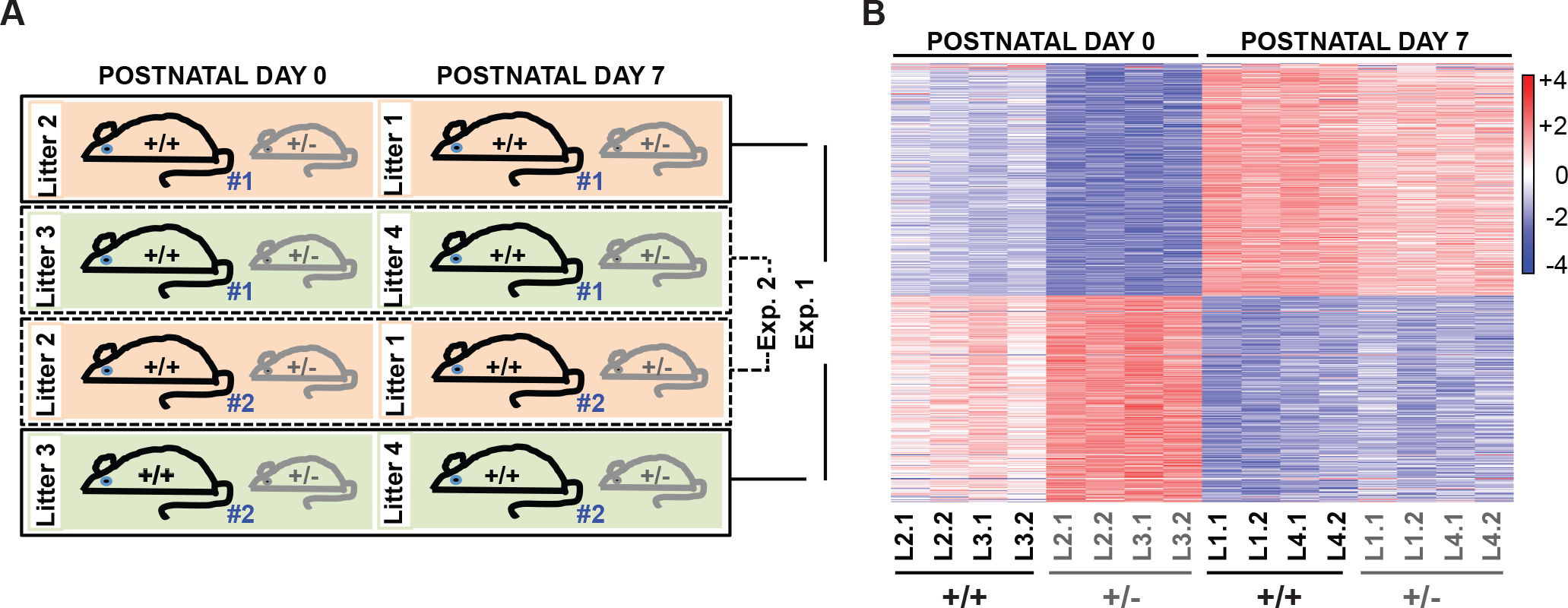
Gene expression in the postnatal *Dyrk1a*^+/+^ and *Dyrk1a*^+/-^cerebral cortex. (A) Experimental design indicating the animal number (#1 or #2), the genotype (*Dyrk1a*^+/+^, +/+; *Dyrk1a*^+/-^, +/-) and the litter (litters 2 and 3 for P0 samples and litters 1 and 4 for P7 samples) used in the Affymetrix array. Experiment (Exp.) 1 and 2 refers to the number of the array cartridge used for hybridization. Two samples for each experimental condition (developmental stage, genotype and litter) were included in each cartridge. (B) Heatmap showing gene expression in WT (*+/+) and Dyrk1a* mutant (+/-) samples at P0 and P7. The rows represent the individual probes and the values in the color-coded scale bar correspond to the standard deviation of each probe from the log2 mean expression value. The differences between the genotypes are smaller than between the developmental stages and they were more evident at P0 than at P7. Note that inter- and intra-litter variability is similar.

**Supplementary Fig. 8.**
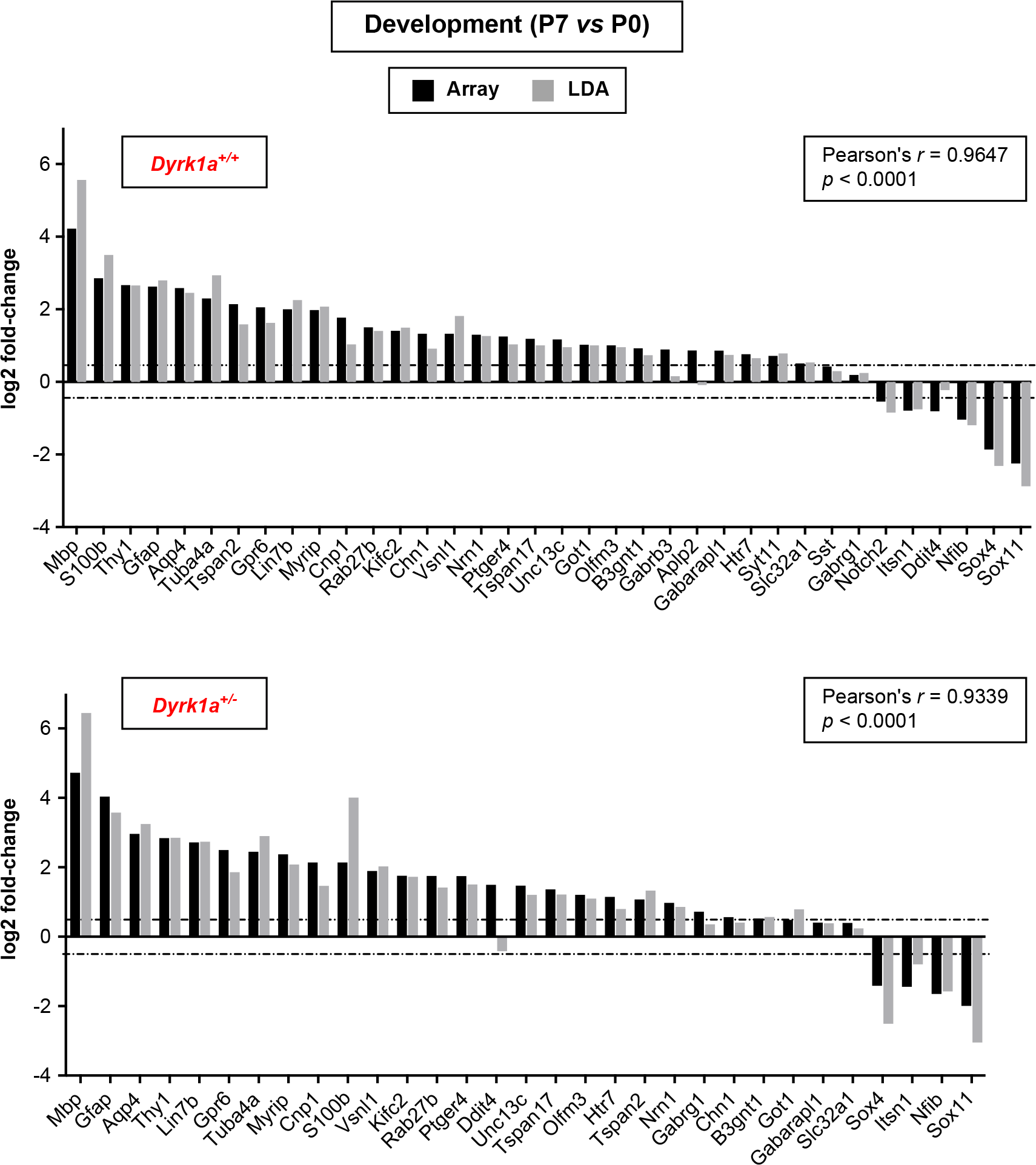
qPCR validation of the microarray data from a low-density array (LDA). Experimental details for the LDA are described in the Supplementary Materials and Methods, and the probes used are listed in Table 5. We selected a set of 36 genes that show the difference in expression between P7 and P0 in the WT (*Dyrk1a*^+/+^) samples on the Affymetrix array (adjusted *p*-value < 0.05). The qPCRs were performed in samples obtained from P0 and P7 *Dyrk1a*^+/+^ and *Dyrk1a*^+/-^ animals (n=4 animals of each genotype and developmental stage). The graphs represent the mean log2 fold-change in value of each gene obtained in the Affymetrix (black bars) and the LDA array (grey bars). Correlation of the log2 fold-change from the microarray and LDA results was calculated with a Pearson’s correlation test. The array and LDA results showed significant correlations in both *Dyrk1a*^+/+^ and *Dyrk1a*^+/-^ samples. The changes in mRNA expression were consistent (up- or down-regulated at P7) through the two experimental approaches, with the exception of *Aplp2* in the *Dyrk1a*^+/+^ samples and *Ddit4* in the *Dyrk1a*^+/-^ samples. The dotted lines indicate the log2 fold-changes equal to ±0.4.

**Supplementary Fig. 9.**
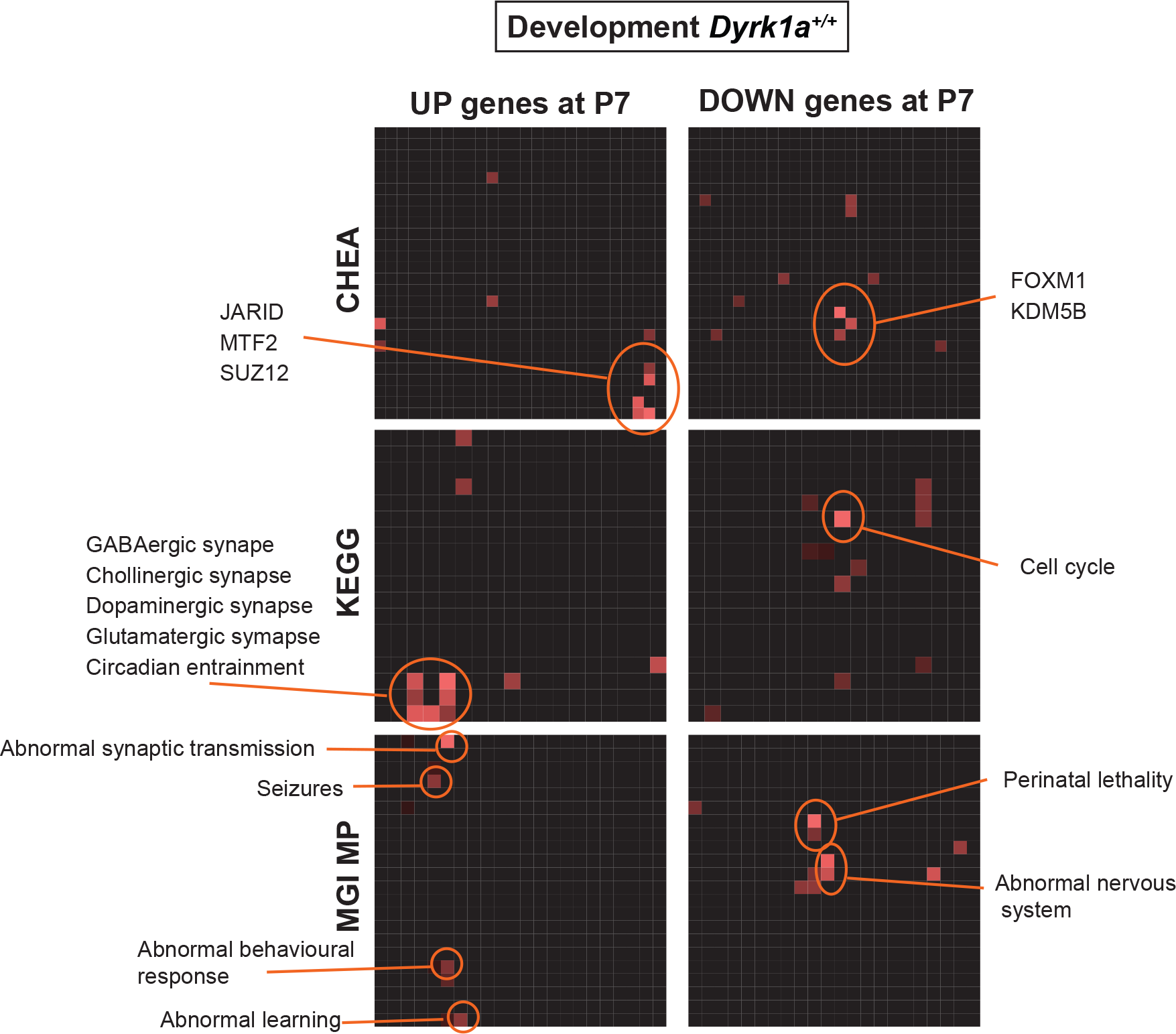
Pathway enrichment analysis of the transcriptome data in *Dyrk1a*^+/+^ cortices during development (P7 vs P0). Enrichment terms for the up-regulated and down-regulated genes at P7 relative to the P0 cortices of WT animals, determined using the Enrichr gene set enrichment analysis and visualization tools. The enriched terms are highlighted in color where the brightness indicates a lower *p*-value. Selective relevant enriched terms are listed on the side on the canvas (ChEA, published ChIP-sequencing data sets; KEGG, cellular and molecular pathways database; MGI-MP, mammalian phenotypes from the Mouse Genome Informatics: see Supplementary Dataset 2 for a complete list of enriched terms). The CHEA analysis indicated that the up-regulated genes were enriched in Polycomb targets, whereas the down-regulated gene set was enriched in targets of the proliferation-associated transcription factor FOXM1. Note that the enrichment analysis of the mammalian phenotype ontology revealed terms like abnormal synaptic transmission, seizures and abnormal learning in the up-regulated gene set, and abnormal nervous system in the down-regulated set.

## Supplementary Movie

Example of a generalized tonic-clonic (GTC) seizure recorded in a *Dyrk1a*^+/-^ mouse. The first manifestation of a GTC seizure is a typical Straub reaction, consisting of a rigid and erected tail lying across the back of the animal in an S-shaped curve. This reaction is followed by facial jerks, myoclonus, the mouse falling on its side and irregular twitches with hyperextension of the limbs, which evolves into a clonic stage with wild jumping and tonic hind-limb extension, and an after-seizure period of immobility. The onset of the GTC seizure is missed in the video-EEG because it occurred while connecting the mouse electrodes to the equipment after placing the mouse in the video-EEG cage.

## References

Alvarez, M., Estivill, X., de la Luna, S., 2003. DYRK1A accumulates in splicing speckles through a novel targeting signal and induces speckle disassembly. J. Cell Sci. 116, 3099–3107.

Aranda, S., Laguna, A., de la Luna, S., 2011. DYRK family of protein kinases: evolutionary relationships, biochemical properties, and functional roles. FASEB J. 25, 449–462.

Arque, G., Fotaki, V., Fernandez, D., Martinez de Lagran, M., Arbones, M.L., Dierssen, M., 2008. Impaired spatial learning strategies and novel object recognition in mice haploinsufficient for the dual specificity tyrosine-regulated kinase-1A (Dyrk1A). PLoS One 3, e2575.

Barallobre, M.J., Perier, C., Bove, J., Laguna, A., Delabar, J.M., Vila, M., Arbones, M.L., 2014. DYRK1A promotes dopaminergic neuron survival in the developing brain and in a mouse model of Parkinson’s disease. Cell Death Dis. 5, e1289.

Becker, W., Soppa, U., Tejedor, F.J., 2014. DYRK1A: a potential drug target for multiple Down syndrome neuropathologies. CNS Neurol. Disord. Drug Targets 13, 26–33.

Benavides-Piccione, R., Dierssen, M., Ballesteros-Yanez, I., Martinez de Lagran, M., Arbones, M.L., Fotaki, V., DeFelipe, J., Elston, G.N., 2005. Alterations in the phenotype of neocortical pyramidal cells in the Dyrk1A+/- mouse. Neurobiol. Dis. 20, 115–122.

Bozzi, Y., Casarosa, S., Caleo, M., 2012. Epilepsy as a neurodevelopmental disorder. Front. Psychiatry 3, 19.

Bronicki, L.M., Redin, C., Drunat, S., Piton, A., Lyons, M., Passemard, S., Baumann, C., Faivre, L., Thevenon, J., Riviere, J.B., et al., 2015. Ten new cases further delineate the syndromic intellectual disability phenotype caused by mutations in DYRK1A. Eur. J. Hum. Genet. 23, 1482–1487.

Courcet, J.B., Faivre, L., Malzac, P., Masurel-Paulet, A., Lopez, E., Callier, P., Lambert, L., Lemesle, M., Thevenon, J., Gigot, N., et al., 2012. The DYRK1A gene is a cause of syndromic intellectual disability with severe microcephaly and epilepsy. J. Med. Genet. 49, 731–736.

Dang, T., Duan, W.Y., Yu, B., Tong, D.L., Cheng, C., Zhang, Y.F., Wu, W., Ye, K., Zhang, W.X., Wu, M., et al., 2018. Autism-associated Dyrk1a truncation mutants impair neuronal dendritic and spine growth and interfere with postnatal cortical development. Mol. Psychiatry 23, 747–758.

de la Torre-Ubieta, L., Won, H., Stein, J.L., Geschwind, D.H., 2016. Advancing the understanding of autism disease mechanisms through genetics. Nat. Med. 22, 345–361.

De Rubeis, S., He, X., Goldberg, A.P., Poultney, C.S., Samocha, K., Cicek, A.E., Kou, Y., Liu, L., Fromer, M., Walker, S., et al., 2014. Synaptic, transcriptional and chromatin genes disrupted in autism. Nature 515, 209–215.

Deciphering Developmental Disorders, S., 2015. Large-scale discovery of novel genetic causes of developmental disorders. Nature 519, 223–228.

Di Vona, C., Bezdan, D., Islam, A.B., Salichs, E., Lopez-Bigas, N., Ossowski, S., de la Luna, S., 2015. Chromatin-wide profiling of DYRK1A reveals a role as a gene-specific RNA polymerase II CTD kinase. Mol. Cell 57, 506–520.

Earl, R.K., Turner, T.N., Mefford, H.C., Hudac, C.M., Gerdts, J., Eichler, E.E., Bernier, R.A., 2017. Clinical phenotype of ASD-associated DYRK1A haploinsufficiency. Mol. Autism 8, 54.

Ernst, C., 2016. Proliferation and differentiation deficits are a major convergence point for neurodevelopmental disorders. Trends Neurosci. 39, 290–299.

Evers, J.M., Laskowski, R.A., Bertolli, M., Clayton-Smith, J., Deshpande, C., Eason, J., Elmslie, F., Flinter, F., Gardiner, C., Hurst, J.A., et al., 2017. Structural analysis of pathogenic mutations in the DYRK1A gene in patients with developmental disorders. Hum. Mol. Genet. 26, 519–526.

Florio, M., Huttner, W.B., 2014. Neural progenitors, neurogenesis and the evolution of the neocortex. Development 141, 2182–2194.

Fotaki, V., Dierssen, M., Alcantara, S., Martinez, S., Marti, E., Casas, C., Visa, J., Soriano, E., Estivill, X., Arbones, M.L., 2002. Dyrk1A haploinsufficiency affects viability and causes developmental delay and abnormal brain morphology in mice. Mol. Cell. Biol. 22, 6636–6647.

Fotaki, V., Martinez De Lagran, M., Estivill, X., Arbones, M., Dierssen, M., 2004. Haploinsufficiency of Dyrk1A in mice leads to specific alterations in the development and regulation of motor activity. Behav. Neurosci. 118, 815–821.

Garcia-Cabrero, A.M., Marinas, A., Guerrero, R., de Cordoba, S.R., Serratosa, J.M., Sanchez, M.P., 2012. Laforin and malin deletions in mice produce similar neurologic impairments. J. Neuropathol. Exp. Neurol. 71, 413–421.

Geschwind, D.H., 2009. Advances in autism. Annu. Rev. Med. 60, 367–380.

Guedj, F., Pereira, P.L., Najas, S., Barallobre, M.J., Chabert, C., Souchet, B., Sebrie, C., Verney, C., Herault, Y., Arbones, M., et al., 2012. DYRK1A: a master regulatory protein controlling brain growth. Neurobiol. Dis. 46, 190–203.

Guimera, J., Casas, C., Pucharcos, C., Solans, A., Domenech, A., Planas, A.M., Ashley, J., Lovett, M., Estivill, X., Pritchard, M.A., 1996. A human homologue of Drosophila minibrain (MNB) is expressed in the neuronal regions affected in Down syndrome and maps to the critical region. Hum. Mol. Genet. 5, 1305–1310.

Himpel, S., Panzer, P., Eirmbter, K., Czajkowska, H., Sayed, M., Packman, L.C., Blundell, T., Kentrup, H., Grotzinger, J., Joost, H.G., et al., 2001. Identification of the autophosphorylation sites and characterization of their effects in the protein kinase DYRK1A. Biochem. J. 359, 497–505.

Himpel, S., Tegge, W., Frank, R., Leder, S., Joost, H.G., Becker, W., 2000. Specificity determinants of substrate recognition by the protein kinase DYRK1A. J. Biol. Chem. 275, 2431–2438.

Huguet, G., Ey, E., Bourgeron, T., 2013. The genetic landscapes of autism spectrum disorders. Annu. Rev. Genomics Hum. Genet. 14, 191–213.

Jabaudon, D., 2017. Fate and freedom in developing neocortical circuits. Nat. Commun. 8, 16042.

Ji, J., Lee, H., Argiropoulos, B., Dorrani, N., Mann, J., Martinez-Agosto, J.A., Gomez-Ospina, N., Gallant, N., Bernstein, J.A., Hudgins, L., et al., 2015. DYRK1A haploinsufficiency causes a new recognizable syndrome with microcephaly, intellectual disability, speech impairment, and distinct facies. Eur. J. Hum. Genet. 23, 1473–1481.

Kannan, N., Neuwald, A.F., 2004. Evolutionary constraints associated with functional specificity of the CMGC protein kinases MAPK, CDK, GSK, SRPK, DYRK, and CK2alpha. Protein Sci. 13, 2059–2077.

Kii, I., Sumida, Y., Goto, T., Sonamoto, R., Okuno, Y., Yoshida, S., Kato-Sumida, T., Koike, Y., Abe, M., Nonaka, Y., et al., 2016. Selective inhibition of the kinase DYRK1A by targeting its folding process. Nat. Commun. 7, 11391.

Kim, O.H., Cho, H.J., Han, E., Hong, T.I., Ariyasiri, K., Choi, J.H., Hwang, K.S., Jeong, Y.M., Yang, S.Y., Yu, K., et al., 2017. Zebrafish knockout of Down syndrome gene, DYRK1A, shows social impairments relevant to autism. Mol. Autism 8, 50.

Kriegstein, A., Alvarez-Buylla, A., 2009. The glial nature of embryonic and adult neural stem cells. Annu. Rev. Neurosci. 32, 149–184.

Lee, E., Lee, J., Kim, E., 2017. Excitation/nhibition imbalance in animal models of autism spectrum disorders. Biol. Psychiatry 81, 838–847.

Li, M., Cui, Z., Niu, Y., Liu, B., Fan, W., Yu, D., Deng, J., 2010. Synaptogenesis in the developing mouse visual cortex. Brain Res. Bull. 81, 107–113.

Lochhead, P.A., Sibbet, G., Morrice, N., Cleghon, V., 2005. Activation-loop autophosphorylation is mediated by a novel transitional intermediate form of DYRKs. Cell 121, 925–936.

Lord, C., Bishop, S.L., 2015. Recent advances in autism research as reflected in DSM-5 criteria for autism spectrum disorder. Annu. Rev. Clin. Psychol. 11, 53–70.

Luco, S.M., Pohl, D., Sell, E., Wagner, J.D., Dyment, D.A., Daoud, H., 2016. Case report of novel DYRK1A mutations in 2 individuals with syndromic intellectual disability and a review of the literature. BMC Med. Genet. 17, 15.

Luttjohann, A., Fabene, P.F., van Luijtelaar, G., 2009. A revised Racine’s scale for PTZ-induced seizures in rats. Physiol. Behav. 98, 579–586.

Markram, H., Toledo-Rodriguez, M., Wang, Y., Gupta, A., Silberberg, G., Wu, C., 2004. Interneurons of the neocortical inhibitory system. Nat. Rev. Neurosci. 5, 793–807.

Martinez de Lagran, M., Benavides-Piccione, R., Ballesteros-Yanez, I., Calvo, M., Morales, M., Fillat, C., Defelipe, J., Ramakers, G.J., Dierssen, M., 2012. Dyrk1A influences neuronal morphogenesis through regulation of cytoskeletal dynamics in mammalian cortical neurons. Cereb. Cortex 22, 2867–2877.

Miller, M.W., 1995. Relationship of the time of origin and death of neurons in rat somatosensory cortex: barrel versus septal cortex and projection versus local circuit neurons. J. Comp. Neurol. 355, 6–14.

Moller, R.S., Kubart, S., Hoeltzenbein, M., Heye, B., Vogel, I., Hansen, C.P., Menzel, C., Ullmann, R., Tommerup, N., Ropers, H.H., et al., 2008. Truncation of the Down syndrome candidate gene DYRK1A in two unrelated patients with microcephaly. Am. J. Hum. Genet. 82, 1165–1170.

Molyneaux, B.J., Arlotta, P., Menezes, J.R., Macklis, J.D., 2007. Neuronal subtype specification in the cerebral cortex. Nat. Rev. Neurosci. 8, 427–437.

Najas, S., Arranz, J., Lochhead, P.A., Ashford, A.L., Oxley, D., Delabar, J.M., Cook, S.J., Barallobre, M.J., Arbones, M.L., 2015. DYRK1A-mediated Cyclin D1 degradation in neural stem cells contributes to the neurogenic cortical defects in Down syndrome. EBioMedicine 2, 120–134.

O’Roak, B.J., Vives, L., Fu, W., Egertson, J.D., Stanaway, I.B., Phelps, I.G., Carvill, G., Kumar, A., Lee, C., Ankenman, K., et al., 2012. Multiplex targeted sequencing identifies recurrently mutated genes in autism spectrum disorders. Science 338, 1619–1622.

Ori-McKenney, K.M., McKenney, R.J., Huang, H.H., Li, T., Meltzer, S., Jan, L.Y., Vale, R.D., Wiita, A.P., Jan, Y.N., 2016. Phosphorylation of beta-tubulin by the Down syndrome kinase, Minibrain/DYRK1A, regulates microtubule dynamics and dendrite morphogenesis. Neuron 90, 551–563.

Packer, A., 2016. Neocortical neurogenesis and the etiology of autism spectrum disorder. Neurosci. Biobehav. Rev. 64, 185–195.

Park, J.H., Jung, M.S., Kim, Y.S., Song, W.J., Chung, S.H., 2012. Phosphorylation of Munc18-1 by Dyrk1A regulates its interaction with Syntaxin 1 and X11alpha. J. Neurochem. 122, 1081–1091.

Posey, J.E., Rosenfeld, J.A., James, R.A., Bainbridge, M., Niu, Z., Wang, X., Dhar, S., Wiszniewski, W., Akdemir, Z.H., Gambin, T., et al., 2016. Molecular diagnostic experience of whole-exome sequencing in adult patients. Genet. Med. 18, 678–685.

Raveau, M., Shimohata, A., Amano, K., Miyamoto, H., Yamakawa, K., 2018. DYRK1A-haploinsufficiency in mice causes autistic-like features and febrile seizures. Neurobiol. Dis. 110, 180–191.

Ruaud, L., Mignot, C., Guet, A., Ohl, C., Nava, C., Heron, D., Keren, B., Depienne, C., Benoit, V., Maystadt, I., et al., 2015. DYRK1A mutations in two unrelated patients. Eur. J. Med. Genet. 58, 168–174.

Rubenstein, J.L., 2010. Three hypotheses for developmental defects that may underlie some forms of autism spectrum disorder. Curr. Opin. Neurol. 23, 118–123.

Somogyi, P., Tamas, G., Lujan, R., Buhl, E.H., 1998. Salient features of synaptic organisation in the cerebral cortex. Brain. Res. Brain Res. Rev. 26, 113–135.

Souchet, B., Guedj, F., Sahun, I., Duchon, A., Daubigney, F., Badel, A., Yanagawa, Y., Barallobre, M.J., Dierssen, M., Yu, E., et al., 2014. Excitation/inhibition balance and learning are modified by Dyrk1a gene dosage. Neurobiol. Dis. 69, 65–75.

Soundararajan, M., Roos, A.K., Savitsky, P., Filippakopoulos, P., Kettenbach, A.N., Olsen, J.V., Gerber, S.A., Eswaran, J., Knapp, S., Elkins, J.M., 2013. Structures of Down syndrome kinases, DYRKs, reveal mechanisms of kinase activation and substrate recognition. Structure 21, 986–996.

Stessman, H.A., Xiong, B., Coe, B.P., Wang, T., Hoekzema, K., Fenckova, M., Kvarnung, M., Gerdts, J., Trinh, S., Cosemans, N., et al., 2017. Targeted sequencing identifies 91 neurodevelopmental-disorder risk genes with autism and developmental-disability biases. Nat. Genet. 49, 515–526.

Sztainberg, Y., Zoghbi, H.Y., 2016. Lessons learned from studying syndromic autism spectrum disorders. Nat. Neurosci. 19, 1408–1417.

Tejedor, F., Zhu, X.R., Kaltenbach, E., Ackermann, A., Baumann, A., Canal, I., Heisenberg, M., Fischbach, K.F., Pongs, O., 1995. minibrain: a new protein kinase family involved in postembryonic neurogenesis in Drosophila. Neuron 14, 287–301.

Tejedor, F.J., Hammerle, B., 2011. MNB/DYRK1A as a multiple regulator of neuronal development. FEBS J. 278, 223–235.

Telley, L., Govindan, S., Prados, J., Stevant, I., Nef, S., Dermitzakis, E., Dayer, A., Jabaudon, D., 2016. Sequential transcriptional waves direct the differentiation of newborn neurons in the mouse neocortex. Science 351, 1443–1446.

Trujillano, D., Bertoli-Avella, A.M., Kumar Kandaswamy, K., Weiss, M.E., Koster, J., Marais, A., Paknia, O., Schroder, R., Garcia-Aznar, J.M., Werber, M., et al., 2017. Clinical exome sequencing: results from 2819 samples reflecting 1000 families. Eur. J. Hum. Genet. 25, 176–182.

van Bon, B.W., Coe, B.P., Bernier, R., Green, C., Gerdts, J., Witherspoon, K., Kleefstra, T., Willemsen, M.H., Kumar, R., Bosco, P., et al., 2016. Disruptive de novo mutations of DYRK1A lead to a syndromic form of autism and ID. Mol. Psychiatry 21, 126–132.

van Bon, B.W., Hoischen, A., Hehir-Kwa, J., de Brouwer, A.P., Ruivenkamp, C., Gijsbers, A.C., Marcelis, C.L., de Leeuw, N., Veltman, J.A., Brunner, H.G., et al., 2011. Intragenic deletion in DYRK1A leads to mental retardation and primary microcephaly. Clin. Genet. 79, 296–299.

Wang, T., Guo, H., Xiong, B., Stessman, H.A., Wu, H., Coe, B.P., Turner, T.N., Liu, Y., Zhao, W., Hoekzema, K., et al., 2016. De novo genic mutations among a Chinese autism spectrum disorder cohort. Nat. Commun. 7, 13316.

Widowati, E.W., Ernst, S., Hausmann, R., Muller-Newen, G., Becker, W., 2018. Functional characterization of DYRK1A missense variants associated with a syndromic form of intellectual deficiency and autism. Biol. Open 7.

Zhang, Y., Kong, W., Gao, Y., Liu, X., Gao, K., Xie, H., Wu, Y., Zhang, Y., Wang, J., Gao, F., et al., 2015. Gene mutation analysis in 253 Chinese children with unexplained epilepsy and intellectual/developmental disabilities. PLoS One 10, e0141782.

## Supplementary References

Benjamini, Y., Hochberg, Y., 1995. Controlling the false discovery rate: a practical and powerful approach to multiple testing. J. R. Stat. Soc. Series B Methodol. 57, 12.

Irizarry, R.A., Hobbs, B., Collin, F., Beazer-Barclay, Y.D., Antonellis, K.J., Scherf, U., Speed, T.P., 2003. Exploration, normalization, and summaries of high density oligonucleotide array probe level data. Biostatistics 4, 249–264.

Kornev, A.P., Haste, N.M., Taylor, S.S., Eyck, L.F., 2006. Surface comparison of active and inactive protein kinases identifies a conserved activation mechanism. Proc. Natl. Acad. Sci. USA 103, 17783–17788.

Kornev, A.P., Taylor, S.S., Ten Eyck, L.F., 2008. A helix scaffold for the assembly of active protein kinases. Proc. Natl. Acad. Sci. USA 105, 14377–14382.

Kuleshov, M.V., Jones, M.R., Rouillard, A.D., Fernandez, N.F., Duan, Q., Wang, Z., Koplev, S., Jenkins, S.L., Jagodnik, K.M., Lachmann, A., et al., 2016. Enrichr: a comprehensive gene set enrichment analysis web server 2016 update. Nucleic Acids Res. 44, W90–97.

Landrum, M.J., Lee, J.M., Benson, M., Brown, G., Chao, C., Chitipiralla, S., Gu, B., Hart, J., Hoffman, D., Hoover, J., et al., 2016. ClinVar: public archive of interpretations of clinically relevant variants. Nucleic Acids Res. 44, D862–868.

Lin, C.W., Hsueh, Y.P., 2014. Sarm1, a neuronal inflammatory regulator, controls social interaction, associative memory and cognitive flexibility in mice. Brain Behav. Immun. 37, 142–151.

Mudunuri, U., Che, A., Yi, M., Stephens, R.M., 2009. bioDBnet: the biological database network. Bioinformatics 25, 555–556.

Paxinos, G., Franklin, K.B.J. (2001). The mouse brain in stereotaxic coordinates, Second edition edn (San Diego, CA.).

Pfaffl, M.W., 2001. A new mathematical model for relative quantification in real-time RT-PCR. Nucleic Acids Res. 29, e45.

Rump, P., Jazayeri, O., van Dijk-Bos, K.K., Johansson, L.F., van Essen, A.J., Verheij, J.B., Veenstra-Knol, H.E., Redeker, E.J., Mannens, M.M., Swertz, M.A., et al., 2016. Whole-exome sequencing is a powerful approach for establishing the etiological diagnosis in patients with intellectual disability and microcephaly. BMC Med. Genomics 9, 7.

Smyth, G.K., 2004. Linear models and empirical bayes methods for assessing differential expression in microarray experiments. Stat. Appl. Genet. Mol. Biol. 3, Article3.

Tan, C.M., Chen, E.Y., Dannenfelser, R., Clark, N.R., Ma’ayan, A., 2013. Network2Canvas: network visualization on a canvas with enrichment analysis. Bioinformatics 29, 1872–1878.

Taylor, S.S., Kornev, A.P., 2011. Protein kinases: evolution of dynamic regulatory proteins. Trends Biochem. Sci. 36, 65–77.

Thomas, A., Burant, A., Bui, N., Graham, D., Yuva-Paylor, L.A., Paylor, R., 2009. Marble burying reflects a repetitive and perseverative behavior more than novelty-induced anxiety. Psychopharmacology (Berl) 204, 361–373.

